# Methylation of histone H4 lysine 20 by PR-Set7 ensures the integrity of late replicating sequence domains in Drosophila

**DOI:** 10.1101/028100

**Authors:** Yulong Li (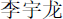), Robin L. Armstrong, Robert J. Duronio, David M. MacAlpine

## Abstract

The methylation state of lysine 20 on histone H4 (H4K20) has been linked to chromatin compaction, transcription, DNA repair and DNA replication. Monomethylation of H4K20 (H4K20me1) is mediated by the cell cycle-regulated histone methyltransferase PR-Set7. PR-Set7 depletion in mammalian cells results in defective S phase progression and the accumulation of DNA damage, which has been partially attributed to defects in origin selection and activation. However, these studies were limited to only a handful of mammalian origins, and it remains unclear how PR-Set7 and H4K20 methylation impact the replication program on a genomic scale. We employed genetic, cytological, and genomic approaches to better understand the role of PR-Set7 and H4K20 methylation in regulating DNA replication and genome stability in *Drosophila* cells. We find that deregulation of H4K20 methylation had no impact on origin activation throughout the genome. Instead, depletion of PR-Set7 and loss of H4K20me1 results in the accumulation of DNA damage and an ATR-dependent cell cycle arrest. Coincident with the ATR-dependent cell cycle arrest, we find increased DNA damage that is specifically limited to late replicating regions of the *Drosophila* genome, suggesting that PR-Set7-mediated monomethylation of H4K20 is critical for maintaining the genomic integrity of late replicating domains.

## INTRODUCTION

Histone post-translational modifications (PTMs) regulate almost all DNA-templated processes including DNA replication, transcription and DNA repair. Deregulation of these epigenetic histone modifications has the potential to lead to catastrophic consequences at both the cellular and organismal level. One such epigenetic mark, methylation of histone H4 lysine 20 (H4K20), is critical for maintaining genome stability, and its deregulation impacts transcription, chromatin compaction, DNA repair, cell cycle progression, and DNA replication [(reviewed in 1, 2, 3)].

Monomethylation of H4K20 (H4K20me1) is catalyzed by the histone methyltransferase PR-Set7/Set8, orthologues of which exist in all metazoans (4, 5). H4K20 can also be di- and tri-methylated by the Suv4-20 h1 and h2 homologs in mammalian cells and a single Suv4-20 in *Drosophila* (6, 7). The levels of mammalian PR-Set7 and H4K20me1 are cell cycle regulated. PR-Set7 harbors a conserved PIP-box motif and undergoes PCNA- and CRL4^Cdt2^-mediated degradation during S phase(8-12). This interaction between the PR-Set7 PIP-box and PCNA is conserved in *Drosophila* cell lines, where the levels of PR-Set7 and H4K20me1 display a similar cell cycle oscillation pattern as seen in mammalian systems(13).

Not only are PR-Set7 and H4K20me1 levels coupled to DNA replication via the PIP-box dependent degradation of PR-Set7, but the DNA replication program is also regulated, in part, by the methylation status of H4K20. Mammalian cells depleted of PR-Set7 are defective in S phase progression, accumulate DNA damage, and activate the DNA damage response (14-16). Mammalian PR-Set7 promotes origin activity at select origins by recruiting pre-Replication Complex (pre-RC) components onto chromatin(11), suggesting that impairment of origin activity in the absence of PR-Set7 may contribute to genome instability. Stabilization of PR-Set7 resulting from the expression of a degradation resistant PIP-box mutant version of PR-Set7 also leads to re-replication and genome instability(11). Similarly, *PR-Set7* mutant *Drosophila* neuroblasts show reduced mitotic and S phase indices(17), and PR-Set7 RNAi treated S2 cells have an increased S phase population(18); however, re-replication resulting from PR-Set7 overexpression has not been observed in the fly.

The ability of mammalian PR-Set7 to regulate replication origin activity is dependent on its catalytic function (11) and the presence of Suv4-20h1/h2, which catalyze the di- and tri-methylation of H4K20(19). Consistent with this, H4K20me2 and H4K20me3 may function to stabilize ORC binding via the BAH domain of ORC1 or the WD40 domain of LRWD1/ORCA (19-21). However, H4K20me2 constitutes more than 80% of total histone H4(7), which implies that 96% of all nucleosomes will contain at least one histone H4 dimethylated at lysine 20. Similarly, trimethylated H4K20 is, for the most part, limited to heterochromatic regions (22, 23). Together, these results suggest that additional mechanisms must exist to specify origin selection in mammalian genomes. Moreover, it is estimated that mammalian cells have more than 40,000 origins of replication(24), while the influence of PR-Set7 on origin licensing has only been examined at a select few origins(11).

Here, we investigate the function of PR-Set7 and H4K20 methylation in regulating the DNA replication program in *Drosophila.* Similar to mammalian systems, loss of PR-Set7 perturbs the normal cell cycle progression of cultured Kc167 cells, presumably due to the accumulation of unmethylated H4K20 over the course of multiple cell cycles. We show that PR-Set7 depletion results in DNA damage and activation of an ATR-dependent cell cycle checkpoint in Kc167 cells, implying a defect in DNA replication. We also observe spontaneous DNA damage in the imaginal wing discs of mutant third instar larvae that have an alanine substitution at position 20 in histone H4 (H4K20A) and are thus unable to undergo methylation on lysine 20(25). Loss of PR-Set7 did not perturb the genome-wide pattern of replication origin activation; instead, we find that late replication domains are specifically sensitive to downregulation of PR-Set7 and accumulate DNA damage in Kc167 cells. Together, these data suggest that PR-Set7 and H4K20me1 are critical for maintaining the integrity of late replicating domains in *Drosophila.*

## MATERIALS AND METHODS

## *Drosophila* cell culture

*Drosophila* Kc167 cells were cultured at 25°C in Schneider's Insect Cell Medium (Invitrogen) supplemented with 10% heat-inactivated fetal bovine serum (Gemini) and 1% penicillin/streptomycin/glutamine (Invitrogen). Dacapo and PR-Set7^PIPm^ were cloned into the pMK33 plasmid under the control of a Cu^2+^ inducible metallothionein promoter and transfected into Kc167 cells with the Effectene Transfection Reagent (Qiagen). Two days after transfection, cells were selected and maintained with 0.125 μg/ml Hygromycin B (Sigma). To arrest cells in G1, Dacapo (26) was overexpressed with 1 mM copper sulfate for 24 hours. Similarly, PR-Set7^PIPm^ was induced with 0.5 mM copper sulfate. To arrest cells at the G1/S transition, 1 mM hydroxyurea (Sigma) was added for 24 hours (unless otherwise noted).

## Double-stranded RNA synthesis and RNA interference

Double-stranded RNAs (dsRNAs) were generated by in vitro transcription (Promega T7 RiboMax) from 400-800 bp PCR products representing the target gene. Target sequences were flanked by T7 promoters on both strands. Primer sequences used are found in Table S2. To perform RNAi, cells were washed and diluted in serum-free medium to a concentration of 2×10^6^ cells/ml, and 15 μg dsRNA per 10^6^ cells were added and incubated for one hour before diluting cells to 1×10^6^ cells/ml with 2x medium. The cells were then incubated at 25°C for 24-72 hours.

## Western blot analysis

Primary antibodies used for western blot analysis: rabbit anti-PR-Set7 (gift from Danny Reinberg) at 1:500, rabbit anti-H4K20me1 (Abcam) at 1:1000, rabbit anti-histone H3 (Abcam) at 1:2000, rabbit anti-Orc6 (gift from Igor Chesnokov) at 1: 1000. Alexa Fluor 680 goat anti-rabbit IgG (Invitrogen) was used at 1:10000 as a secondary antibody. Immunoblots were scanned using a LiCor infrared system and analyzed with Odyssey software.

## Flow cytometry and cell sorting

For cell cycle analysis by flow cytometry, cells were harvested by centrifugation, washed in 1x cold PBS, resuspended in 500 μl PBS and fixed with 5 ml ice-cold 100% ethanol at 4°C overnight. The cells were then washed in 1x cold PBS, resuspended in PBS with 0.3 mg/ml RNase A and 15 μg/ml propidium iodide, and incubated for 30 minutes at room temperature. Flow cytometry was performed on BD FACscan analyzer machine and 30000 cells were recorded for each sample. Histograms of cell cycle distribution were generated with the R bioconductor flowCore package(27).

To sort cells into G1/S and late S/G2 populations, asynchronous cells were fixed with 1% formaldehyde for 10 min and quenched by 125 mM glycine for 10 min. Then 0.5 μg/ml Hoechst 33342 (Invitrogen) was used to stain DNA before the cells were sorted into G1/S and late S/G2 populations based on DNA content using BD DiVa sorter.

## Immunofluorescence

For EdU pulse labeling, cells were incubated with 10 μM EdU for 30 minutes before harvest, fixed with 4% paraformaldehyde in PBS for 10 minutes, permeabilized with 0.5% Triton X-100 in PBS for 3 minutes, and labeled with the Click-iT EdU imaging kit (Invitrogen). Cells were then blocked with 3% BSA in PBS with 0.1% Triton X-100 (PBS-T) and stained with rabbit anti-H4K20me1 (Abcam) at a dilution of 1:1000. Alexa Fluor 568 goat anti-rabbit (Invitrogen) was used at 1:500 as the secondary antibody. For detecting γ-H2A.v, cells were fixed, permeabilized, and blocked as above. Cells were stained with mouse monoclonal anti-γ-H2A.v (DSHB UNC93-5.2.1-s) at 1:500 and Alexa Fluor 488 goat anti-mouse (Invitrogen) at 1:500. All slides were mounted with VECTASHIELD Mounting Medium with DAPI (Vector Laboratories) and observed with Zeiss Axio Imager widefield fluorescence microscope with 40x objective.

For γ-H2A.v staining of wing discs, third instar wandering larval cuticles were inverted and fixed in 3.7% paraformaldehyde in PBS for 25 minutes at 25°C. Cuticles were washed for 1 hour in PBS-T, and then blocked in PBS with 5% NGS for 1 hour at 25°C. Mouse anti-γ-H2A.v primary antibody (28) was used at 1: 10000 for 16 hours at 4°C. Cy5 conjugated goat α-mouse was used at 1:1000 for 2 hours at 25°C. DNA was counterstained with DAPI, and the discs were mounted in ProLong Gold antifade reagent. Images were captured using a Leica TCS SP5.

## DNA comet assay

The neutral DNA comet assay was performed using the CometAssay Kit (Trevigen). DNA was stained with the SYTOX Green nucleic acid stain (Invitrogen) diluted to 1:10000 in TE and images were acquired with a Zeiss Axio Imager widefield fluorescence microscope. The comet tail moment was quantified using the ImageJ plugin OpenComet(29).

## Chromatin immunoprecipitation

Chromatin immunoprecipitations were performed as previously described(30). H4K20me1 and γ-H2A.v ChIP experiments was performed using rabbit anti-H4K20me1 (Abcam, validated for ChIP-seq(31)) and mouse anti-γ-H2A.v (DSHB UNC93-5.2.1-s) antibodies, respectively. The γ-H2A.v ChIP experiment was performed in duplicate. The H4K20me1 experiment was also performed in duplicate by using two independent approaches (HU exposure and cell sorting) to obtain synchronous cell populations.

## Genome-wide analysis of early origin activation

A 10 ml culture of asynchronous cells at a concentration of 1×10^6^ cells/ml was treated with dsRNA or copper for 36 hours and then 1 mM HU was added and the cells were incubated for 4 hours to stall any active replication forks. Subsequently, 10 μM BrdU was added to the culture and the cells were incubated for another 16 hours before harvesting. Genomic DNA extraction and BrdU immunoprecipitation were performed as previously described(32).

## Sequencing library preparation

Illumina sequencing libraries were generated from the DNA enriched by ChIP or BrdU immunoprecipitation using Illumina's TruSeq protocols.

## Sequencing read alignment

Sequenced reads were aligned to the dm3 release 5.12 of the *Drosophila melanogaster* genome assembly (downloaded from the UCSC Genome Browser, and available at ftp://hgdownload.cse.ucsc.edu/goldenPath/dm3/bigZips/chromFa.tar.gz), using the software package Bowtie (version 0.12.7)(33). The following Bowtie parameters were used: -n 2 -l 30 -m 1 --best --strata -y.

## Data accession and genomic data analysis

All genomic data are publicly available at the NCBI GEO data repository with the accession number GSE73668.

For H4K20me1 ChIP-seq and BrdU-seq experiments, RPKM (reads per kilobase, per million mappable reads) was calculated over bins of 10 kb stepping every 1 kb. For the γ-H2A.v datasets, RPKM was determined over nonoverlapping bins of 5 kb. The pearson correlation between individual biological replicates for each experiment are listed in Table S3. The mean RPKM of replicate datasets were used for analysis. The list of gene regions was downloaded from UCSC Table Browser (Assembly BDGP R5/dm3).

The H4K20me1 ChIP signal was normalized between early and late S phase samples based on the relative enrichment in the top 5% of detected peaks(34).

For γ-H2A.v ChIP-seq analysis, we calculated the log2 difference of ChIP signal: log2 difference = log2(PR-Set7 RNAi RPKM + 1) - log2(control RNAi RPKM +1). A positive value indicates increased γ-H2A.v upon PR-Set7 RNAi treatment. To identify discrete γ-H2A.v domains, we trained a two-state hidden Markov model (HMM, representing positive or negative γ-H2A.v domains), with each state emitting a univariate normal distribution corresponding to the log2 difference values. To generate the metaplot of replication timing over late replicating domains, each late domain was divided into 50 bins. Twenty-five flanking upstream and downstream bins, equivalent in width to the late domain bin, were also included. The γ-H2A.v log2 differences and replication timing values within each bin were then calculated and the respective means across all domains were plotted to generate a metadomain.

## RESULTS

## H4K20 monomethylation is critical for the completion of the subsequent cell cycle

The methylation state (none, mono-, di- or tri-methylation) of H4K20 has been linked to genome stability and cell cycle progression in a number of model systems(reviewed in 1). To assess the impact of H4K20 methylation states on cell cycle progression in *Drosophila* Kc167 cells, we manipulated the levels of the H4K20 methyltransferases, PR-Set7 and Suv4-20 (Fig 1A). Depletion of the monomethyltransferase PR-Set7 with double-stranded RNA (dsRNA) mediated RNAi for 72 hours resulted in a reduction of H4K20me1 levels compared with control RNAi. Conversely, depletion of Suv4-20, which catalyzes di- and tri-methylation of H4K20, resulted in a dramatic increase in H4K20me1 levels. We also observed a slight increase in H4K20me1 levels relative to control RNAi when simultaneously depleting PR-Set7 and Suv4-20. This slight increase is likely due to differences between the rates of H4K20 monomethylation by PR-Set7 and subsequent dimethylation by Suv4-20. Overexpression of a stabilized PIP box mutant of PR-Set7 (F523A and F524A; PR-Set7^PIPm^) under the control of a copper-inducible metallothionein promoter did not increase the levels of H4K20me1 compared with control RNAi, presumably due to the rapid conversion of H4K20me1 to higher states of methylation(8). We only observed a very modest increase in PR-Set7 levels after overexpression of PR-Set7^PIPm^ in asynchronous cells (Fig 1A). We hypothesize that multiple cell cycle-dependent mechanisms contribute to the regulation of *Drosophila* PR-Set7 levels, as we observed significant overexpression of PR-Set7^PIPm^ in hydroxyurea (HU) treated cells that were arrested at the G1/S transition (Fig S1A).

**Figure 1.**
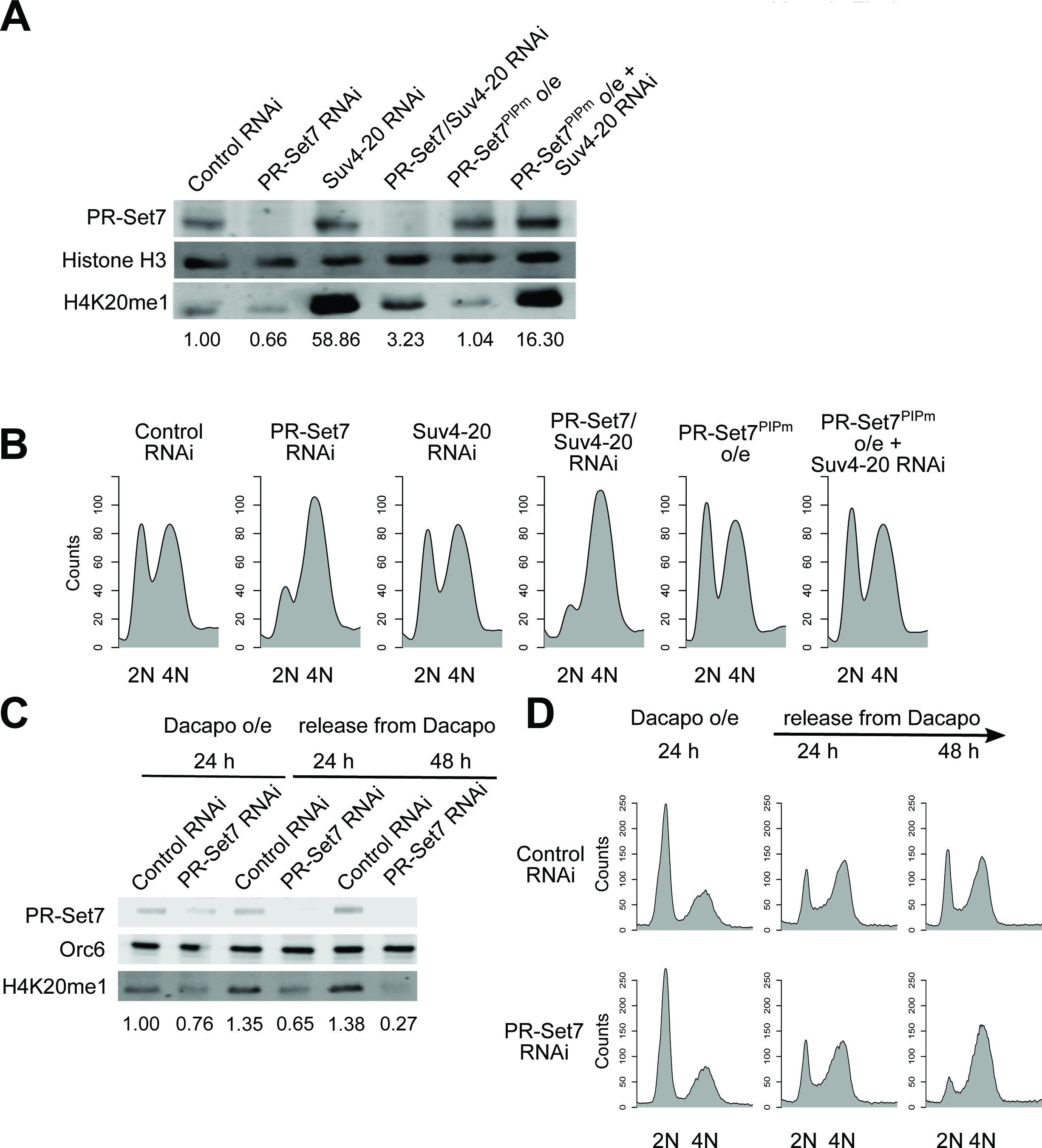
PR-Set7 is necessary for cell cycle progression. (A-B) PR-Set7 downregulation arrests cell cycle progression. Asynchronous cells were depleted for the histone methyltransferases, PR-Set7 or Suv4-20, by RNAi, or alternatively, a stabilized mutant of PR-Set7, PR-Set7^PIPm^, was overexpressed alone or in combination with Suv4-20 depletion for 72 hours. Cells were harvested for (A) western blot analysis for relative levels of PR-Set7 and H4K20me1 (histone H3 as the loading control), and for (B) cell cycle analysis by flow cytometry. (C-D) Kinetics of PR-Set7 RNAi induced cell cycle arrest. Dacapo was overexpressed for 24 hours with control or PR-Set7 dsRNA. The cells were then released from Dacapo overexpression and collected at 24 hours and 48 hours for (C) western blot analysis of PR-Set7 and H4K20me1 (Orc6 as the loading control) and (D) flow cytometry analysis of the cell cycle profile.

Cell cycle progression was monitored in parallel by flow cytometry for each of the genetic manipulations described above (Fig 1B). Depletion of PR-Set7 for 72 hours in Kc167 cells resulted in an accumulation of cells with 4N or near 4N DNA content, similar to previous reports in *Drosophila* S2 cells(13). Interestingly, we also observed an accumulation of cells with 4N DNA content following the simultaneous depletion of PR-Set7 and Suv4-20 despite the slight increase in H4K20me1. Given that PR-Set7 activity is strictly coupled to the cell cycle and H4K20me1 is a transient epigenetic mark deposited in late S phase and converted to H4K20me2/3 by late mitosis(8, 13), we hypothesize that the cell cycle arrest may not be due to the depletion of parental H4K20me1 levels, but rather the accumulation of nascent histone H4.

To test this hypothesis, we examined the kinetics of the cell cycle arrest following PR-Set7 downregulation in synchronized cells (Figs 1C and 1D). Cells were synchronized in G1 by overexpressing the *Drosophila* p27 homolog, Dacapo(35), in the presence of control or PR-Set7 dsRNA. The cells were then released from the Dacapo-induced arrest and allowed to re-enter the cell cycle. In contrast to control cells, cells depleted for PR-Set7 exhibited no increase in H4K20me1 following release from the Dacapo-induced G1 cell cycle arrest (Fig S2). Despite the absence of PR-Set7 and a failure to monomethylate H4K20 during S phase, the cell cycle profile for PR-Set7 RNAi treated cells was indistinguishable from control RNAi treated cells after 24 hours, implying a decoupling of cell cycle progression and H4K20me1 levels. Only after 48 hours did we observe a strong cell cycle arrest. We propose that the monomethylation of nascent H4K20 deposited during S phase is critical for the completion of the next cell cycle.

## Global H4K20 monomethylation occurs in late S phase

Cell cycle regulated levels of PR-Set7 start to peak in late S phase with the accumulation of H4K20 monomethylation(8, 13). Recent reports indicate that *Drosophila* PR-Set7 is coupled to the replication fork via an interaction with DNA Pol alpha(13), which may suggest that the accumulation of H4K20 monomethylation in late S phase will be associated with recently replicated sequences. We used cytological and genomic approaches to characterize and monitor the accumulation of H4K20 monomethylation throughout the cell cycle in *Drosophila* Kc167 cells. We first assessed cell cycle position and H4K20 monomethylation at the single cell level in an asynchronous cell population by immunofluorescence (Figs 2A and 2B). The incorporation and distribution of the nucleoside analog, 5-ethynyl-2’-deoxyuridine (EdU), served as a proxy for cell cycle position. Cells lacking EdU were assumed to be in G1 or G2, whereas cells in S phase and undergoing DNA replication were labeled with EdU. *Drosophila* cells in early or late S phase are readily differentiated by the pattern of EdU incorporation throughout the nucleus(36). Cells in early S phase exhibit EdU incorporation throughout the majority of the nucleus, with the exception of the pericentric heterochromatin which is easily visualized as a "DAPI bright" nuclear focus. In contrast, EdU incorporation is limited to the pericentric heterochromatin during late S phase. We found that cells in early S phase had very low levels of H4K20me1 staining. In contrast, cells in late S phase exhibited significantly elevated H4K20me1 staining throughout the nucleus. Importantly, the increase in H4K20me1 staining was not restricted to the late replicating pericentric heterochromatin, suggesting that the majority of H4K20 monomethylation is decoupled from the replication fork and active DNA synthesis.

To more precisely ascertain the patterns of H4K20 monomethylation, we used chromatin immunoprecipitation coupled with high-throughput sequencing (ChIP-seq) to map H4K20 monomethylation at the beginning or end of S phase. Cells were synchronized at the G1/S transition (early S phase) by treatment with hydroxyurea (HU), an inhibitor of ribonucleotide reductase. To obtain a synchronous population of late S phase cells, cells were released from the HU arrest and allowed to progress through S phase for six hours (Fig 2C).

**Figure 2.**
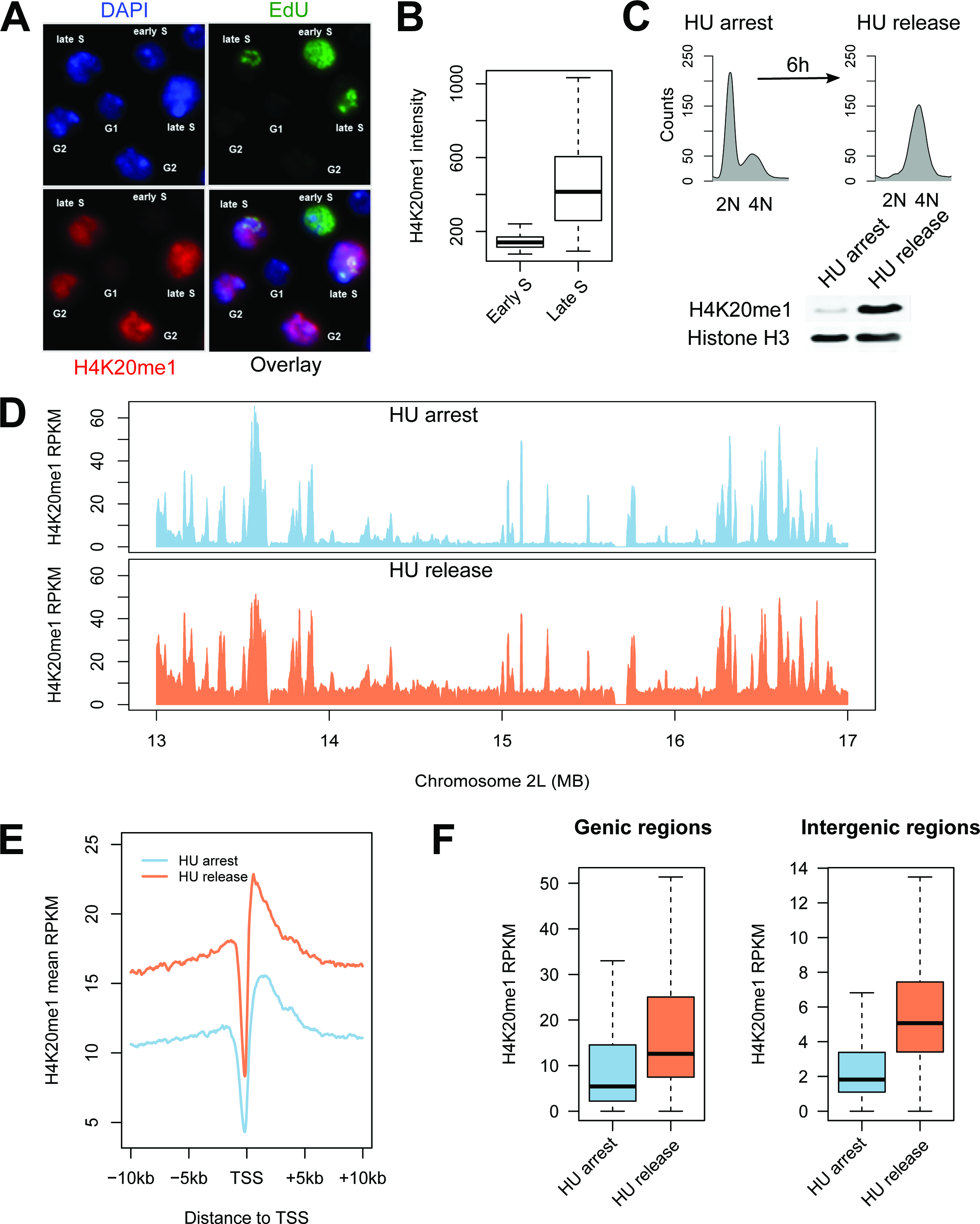
H4K20me1 levels are globally increased throughout the genome in late S phase. (A) H4K20me1 is restricted to cells in late S phase and G2. Asynchronous cells were pulse labeled with EdU for 30 minutes to detect cells in early/mid and late S phase. EdU and H4K20me1 were detected by immunofluorescence microscopy and the nuclei were stained with DAPI. (B) Quantification of H4K20me1 immunofluorescence intensity in early and late S phase (n>20). (C) H4K20me1 is established in late S phase cells released (6 hours) from an HU arrest. (D) Enrichment of H4K20me1 (RPKM) over a representative 4 MB region of chromosome 2L for cells arrested at the G1/S transition with HU (blue) or cells in late S phase following release from HU for 6 hours (orange). (E) H4K20me1 is enriched at transcription start sites and gene bodies. Aggregate plots of H4K20me1 enrichment in HU arrested (blue) and HU released (orange) cells relative to transcription start sites. (F) H4K20me1 levels increase in both genic and intergenic regions during late S phase. Boxplots representing the distribution of H4K20me1 RPKM within 13428 genic and 11430 intergenic regions (p<2.2×10^−16^).

H4K20me1 levels are cell cycle regulated and exhibit their lowest levels in early S phase. Despite only detecting background levels of H4K20me1 by western blot in cells arrested in early S phase at the G1/S transition (Fig 2C), we identified discrete peaks of H4K20me1 enrichment throughout the genome (Fig 2D). These peaks coincided with transcription start sites and regulatory elements as previously noted (Fig 2E;(37)). In late S phase, in addition to enrichment of H4K20me1 at regulatory elements (Fig 2D), we also observed a global increase in H4K20me1 in both genic and intergenic sequences throughout the *Drosophila* genome (Fig 2F). As an alternative approach to isolate cell populations in early or late S phase, we used cell sorting. Chromatin was isolated from these distinct cell populations and the genome-wide distribution of H4K20me1 was determined for early and late S phase. The results were similar to the HU treated samples (Fig S3). Together, these results indicate that monomethylation of H4K20 by PR-Set7 is not limited to replication forks but occurs throughout the genome in late S phase.

## Loss of *Drosophila* PR-Set7 induces DNA damage and activates the ATR checkpoint

Loss of PR-Set7 results in an altered cell cycle distribution with an accumulation of cells with greater than 2N DNA (Fig 1B;(13, 15)), suggesting the activation of the DNA damage response. Prior experiments investigating the occurrence of PR-Set7-mediated DNA damage by immunostaining of the phosphorylated histone H2A.v (γ-H2A.v, the *Drosophila* homolog of H2AX) led to conflicting results(13, 17, 18). The discrepancy in whether or not DNA damage accumulates in PR-Set7 depleted or mutant cells may, in part, result from the specificity of the antibodies used to detect γ-H2A.v. Using a recently developed monoclonal anti-γ-H2A.v antibody(28), we observed a significant increase in the percentage of γ-H2A.v positive cells when Kc167 cells were depleted of PR-Set7 by RNAi treatment for 72 hours (Figs 3A and 3B). The observed increase in γ-H2A.v resulting from PR-Set7 depletion was supported by the direct detection of double-strand DNA breaks (DSBs) using a neutral DNA comet assay (Figs 3C and 3D).

**Figure 3.**
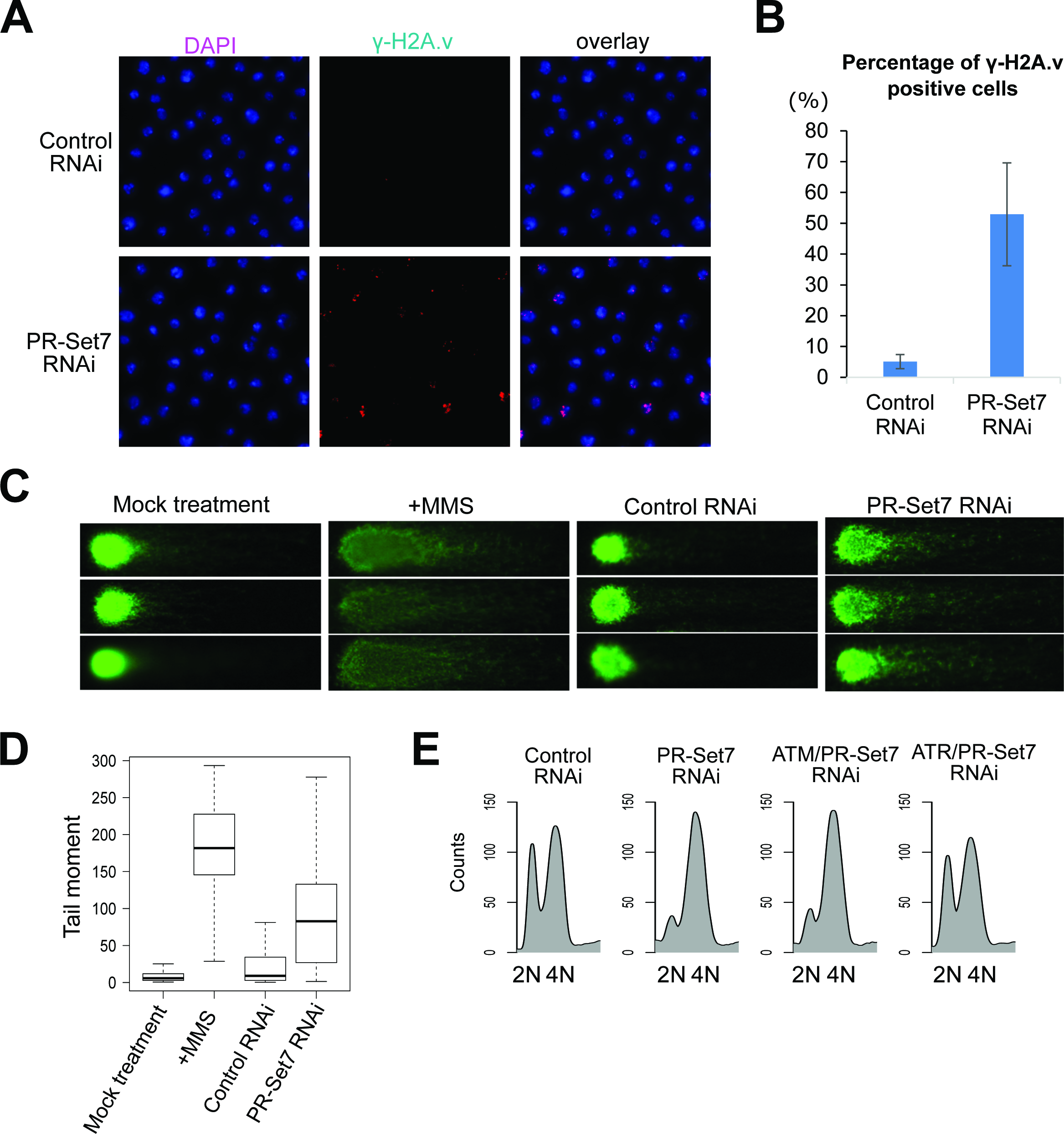
PR-Set7 maintains genome integrity. (A) Depletion of PR-Set7 by RNAi treatment for 72 hours leads to elevated γ-H2A.v staining (red). Nuclear DNA is stained with DAPI (blue). (B) Quantification of the percentage of γ-H2A.v positive cells shown in (A) (mean±SD; n = 3 biological replicates with more than 550 cells per replicate). (C) Depletion of PR-Set7 results in DNA double-strand breaks as determined by a neutral DNA comet assay. Treatment with 0.03% MMS for 6 hours serves as a positive control. (D) Quantification of tail moments (tail length × percentage of DNA in tail) from the neutral DNA comet assay (n>45 for each condition). (E) Flow cytometry analysis of the cell cycle after RNAi treatment against control, PR-Set7, ATM/PR-Set7 and ATR/PR-Set7 for 72 hours.

The accumulation of DNA damage in the absence of PR-Set7 activity led us to test the role of ATM and ATR in mediating the cell cycle arrest. As previously reported(17), we found that depletion of ATR abrogated the checkpoint and cell cycle arrest due to loss of PR-Set7. In contrast to the co-depletion of ATR and PR-Set7, loss of ATM and PR-Set7 had no impact on the cell cycle arrest (Fig 3E). To confirm the efficacy of ATM RNAi, we tested its ability to reduce the phosphorylation of H2A.v when cells were stressed with methyl methanesulfonate (MMS), a strong DNA damaging agent. While ATR or ATM dsRNA alone did not reduce the levels of γ-H2A.v detected by immunofluorescence, the downregulation of both significantly reduced the H2A.v phosphorylation that occurred in response to MMS (Fig S4). We propose that PR-Set7 depletion in *Drosophila* Kc167 cells results in aberrant DNA replication which activates the ATR-dependent intra-S phase checkpoint.

Identified targets of PR-Set7’s catalytic activity include both H4K20 and nonhistone proteins such as p53 (38) and PCNA(39), both of which are critical cell cycle regulators. Thus, it remains possible that PR-Set7’s impact on cell cycle progression and genome integrity may be due to non-histone targets. Indeed, more than 50% of mutant flies harboring an alanine substitution at lysine 20 on histone H4 (H4K20A) were viable(25), which is in stark contrast with PR-Set7 null flies which die at the larval/pupal transition(40). To determine whether loss of H4K20 methylation in the absence of PR-Set7 contributes to the maintenance of genome integrity, we examined H4K20A mutant larval wing discs for the occurrence of increased DNA damage. Relative to control larvae, we observed a 4-fold increase of γ-H2A.v foci in the H4K20A mutant larval wing discs (Fig 4), suggesting that methylation of H4K20 serves as a bona fide PR-Set7 target to maintain genome integrity in *Drosophila.*

**Figure 4.**
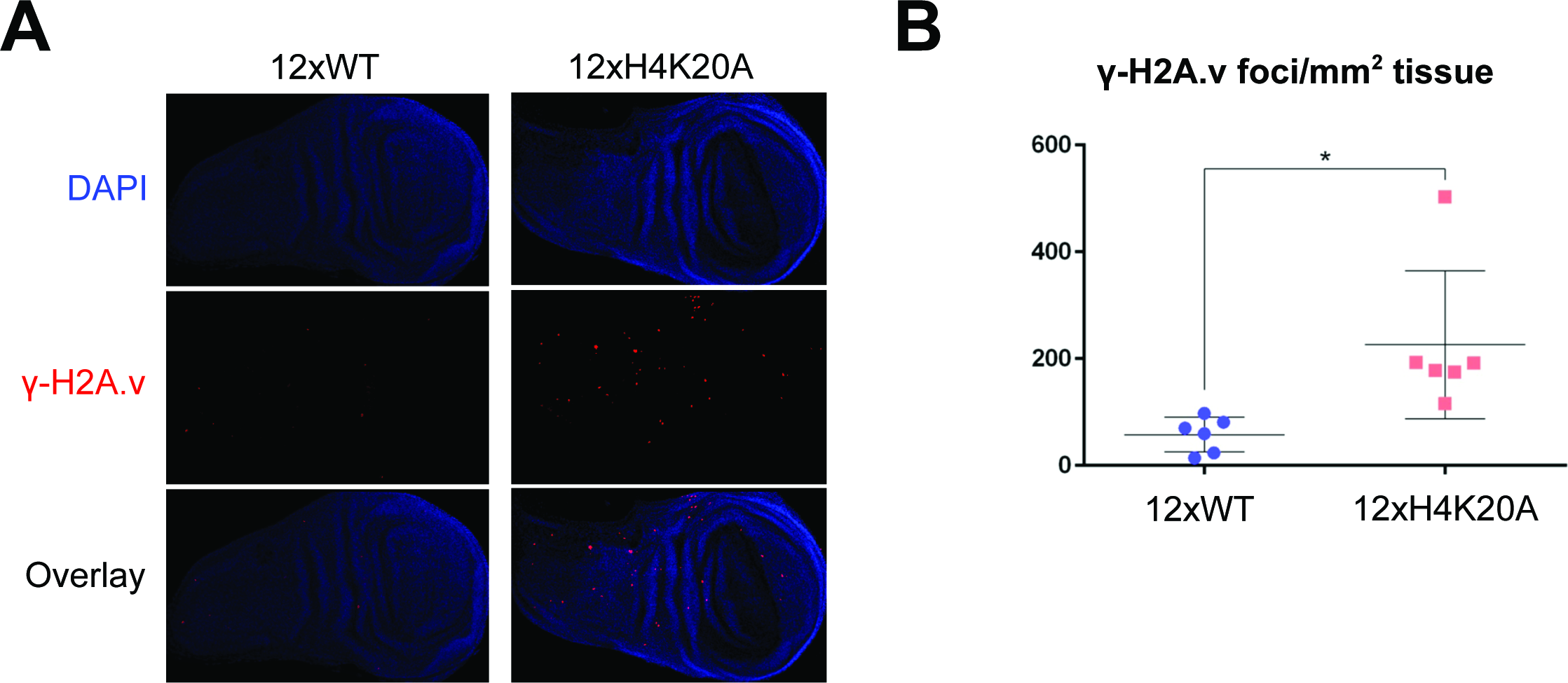
Figure 4. H4K20me1 maintains genome integrity in larval wing discs. (A) Wildtype (WT) control or H4K20A mutant third instar larval imaginal wing discs are stained for γ-H2A.v (red). Nuclear DNA is stained with DAPI (blue). (B) Quantification of γ-H2A.v foci number per mm^2^ tissue (n = 6 imaginal wing discs per genotype; p = 0.016, t-test).

## *Drosophila* H4K20 methylation states do not affect replication origin activity

In mammalian systems, PR-Set7 is thought to regulate pre-RC assembly and origin activation (11), presumably dependent on Suv4-20h1/h2 mediated di- or trimethylation of H4K20(19). We first hypothesized that loss of PR-Set7 may activate the ATR-dependent intra-S phase checkpoint due to aberrant (reduced) origin activation. We used BrdU-seq to map the genome-wide origin activation profile(41). Briefly, we incubated asynchronous cells with dsRNA targeting PR-Set7 or a control sequence for 36 hours before adding HU for 4 hours to inhibit active replication forks. We then added BrdU (in the presence of HU) for another 16 hours. This provided the cells sufficient time to enter the second cell cycle following RNAi treatment so that most cells would be arrested at the second G1/S transition, and only those sequences proximal to early activating origins would incorporate BrdU. As expected, the BrdU incorporation observed in both the control and PR-Set7 depleted cells peaked at ORC binding sites (Fig S5). However, we found no statistical difference in early origin activation between control or PR-Set7 depleted cells (Figs 5A and 5B).

**Figure 5.**
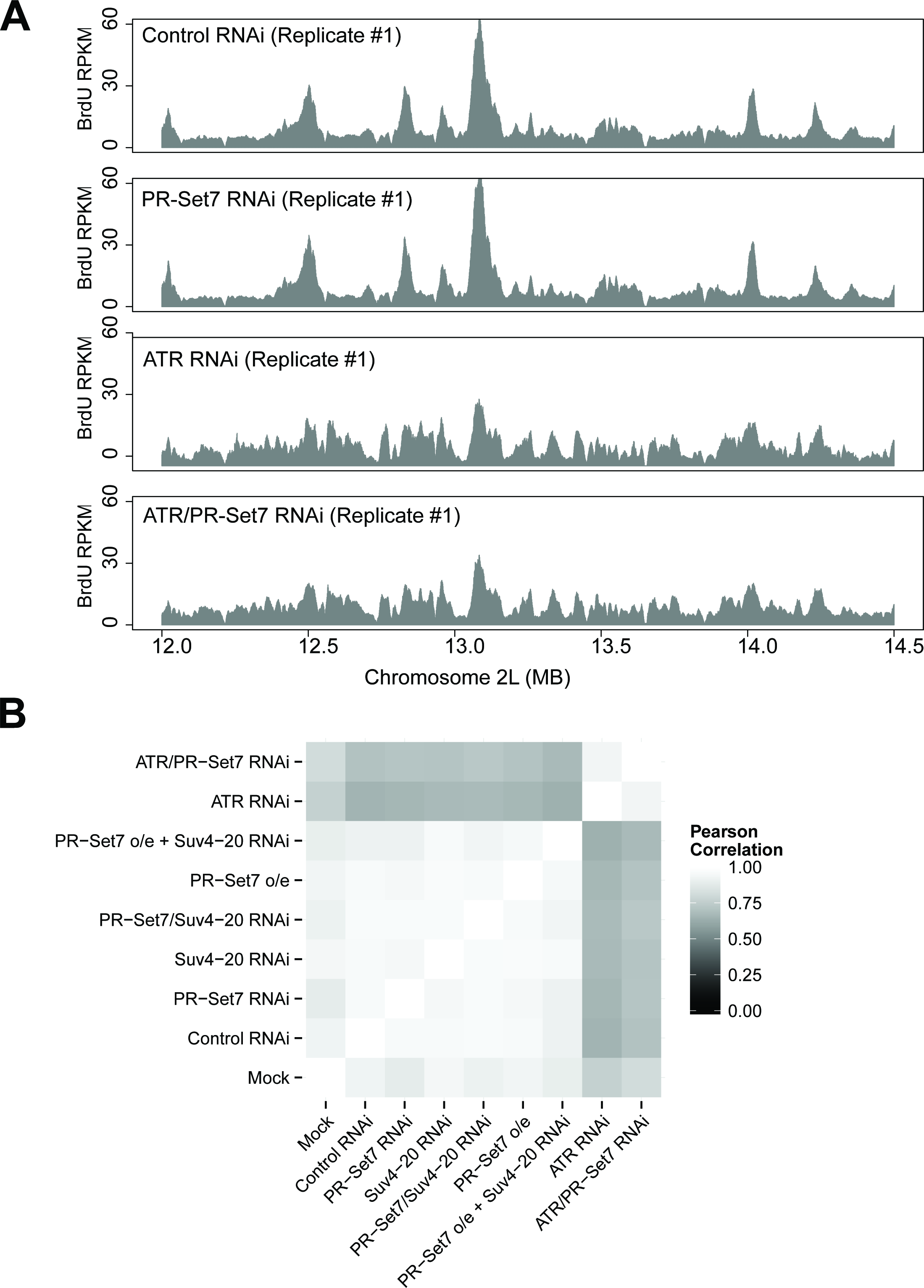
Figure 5. De-regulation of H4K20 methylation does not impact the activation of replication origins. (A) Genome-wide early origin activity was mapped by BrdU-Seq for control and PR-Set7 depleted cells. Late origin activity was mapped by BrdU-Seq for cells depleted of ATR and ATR/PR-Set7. BrdU enrichment (RPKM) is depicted for a representative 2.5 MB window of chromosome 2L. (B) Heatmap depicting the pairwise Pearson correlations for origin activity across the different experimental conditions.

We next considered the hypothesis that PR-Set7 depletion might impair late origin activation. To test this hypothesis, we co-depleted PR-Set7 and ATR in the presence of HU and BrdU. ATR depletion and abrogation of the intra-S phase checkpoint permitted the activation of late and/or dormant origins in addition to early origins. However, the origin activation profile upon co-depletion of ATR and PR-Set7 was indistinguishable from the depletion of ATR alone (Figs 5A and 5B). These results suggest that the cell cycle arrest following depletion of PR-Set7 is not due to a defect in origin activation.

Finally, deregulation of PR-Set7 by overexpression results in DNA re-replication and DNA damage in mammalian cells(11). However, *Drosophila* PR-Set7^PIPm^ overexpression did not lead to re-replication, nor did Suv4-20 RNAi result in any cell cycle defects (Fig 1B). To test if any of these conditions might impact origin activation while keeping the cell cycle progression intact, we performed BrdU-seq in the presence of HU as described above. We detected no significant perturbations in the distribution or activity of replication origins throughout the genome (Fig 5B).

We conclude that deregulation of H4K20 methylation does not directly impact replication origin activity in *Drosophila.*

## Loss of PR-Set7 accumulates DNA damage in late replicating domains

Since replication origin activity was not impacted by PR-Set7 depletion, we hypothesized that the DNA damage lesions (Fig 3) may occur during replicative stress and result from collapsing replication forks. The presence of DNA breaks marked by γ-H2A.v provided an opportunity to identify the genomic locations and features associated with DNA damage resulting from the loss of PR-Set7. We used ChIP-Seq to identify the genome-wide distribution of γ-H2A.v. The control and PR-Set7 depleted samples exhibited numerous γ-H2A.v peaks (Fig S6), presumably reflecting endogenous replication- and transcription-mediated stress(42). In order to identify the genomic locations specifically enriched for γ-H2A.v by PR-Set7 depletion, we normalized the γ-H2A.v signal by calculating the log2 difference in signal between PR-Set7 and control RNAi treated samples (see Materials and Methods). Following normalization, we observed discrete domains of elevated γ-H2A.v throughout the genome (Fig 6A). The vast majority of γ-H2A.v enriched domains were associated with euchromatic late replicating sequences and were inversely correlated with replication timing data (Pearson correlation = -0.696; Figs 6A and 6B; (43)). We used a hidden Markov model to define 102 γ-H2A.v domains and compared them with the replication timing domains identified in a prior study(43). We found that 88 of the γ-H2A.v domains overlapped with a late replicating domain while 120 of the 166 late replicating domains coincided with a γ-H2A.v domain (Table S1). We did not observe hotspots of γ-H2A.v within the late domains, but rather found that γ-H2A.v spanned the entire late domain region (Fig 6C). Given that ChIP-seq is a population-based assay averaged over millions of cells, the lack of discrete γ-H2A.v ‘hotspot’ peaks within the late replicating domains implies a spatial pattern of stochastic DNA damage likely resulting from replication fork collapse in the absence of H4K20me1.

**Figure 6.**
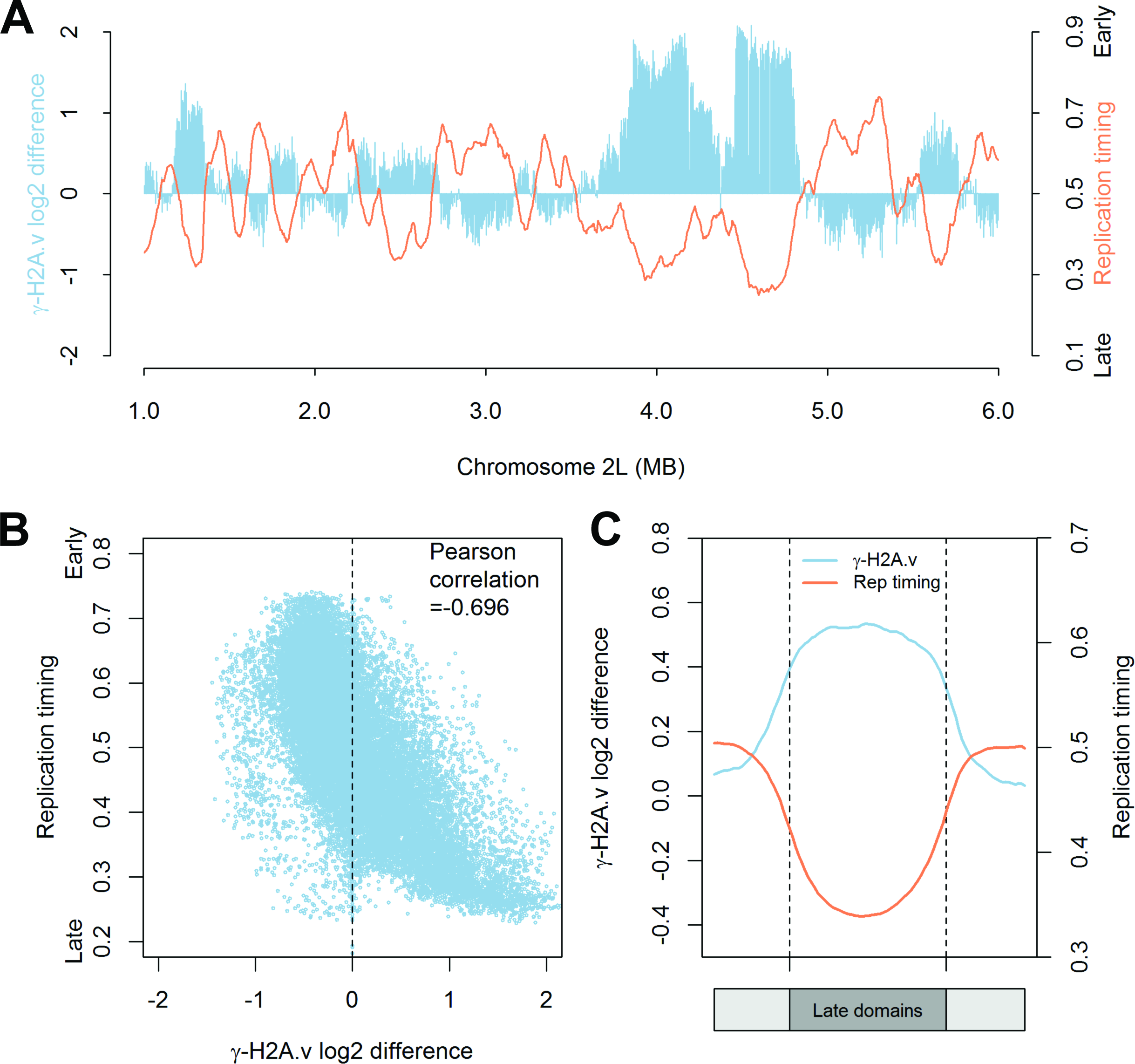
Loss of PR-Set7 leads to the accumulation of DNA damage in late replicating regions of the *Drosophila* genome. (A) The DNA damage marker, γ-H2A.v, accumulates in late replicating regions of the genome. The genome-wide distribution of γ-H2A.v was determined for control and PR-Set7 depleted cells by ChIP-Seq (Fig S6). The log2 difference in γ-H2A.v enrichment between PR-Set7 and control RNAi treated cells (blue) is depicted for a 5 Mb window of chromosome 2L. A replication timing profile (orange) (43) for chromosome 2L is overlaid on the plot. (B) γ-H2A.v is inversely correlated with the time of replication in S phase. Each data point represents a 5 kb bin and the Pearson correlation between replication timing and the log2 difference in γ-H2A.v between PR-Set7 depleted and control cells was calculated. (C) γ-H2A.v is enriched throughout late replicating domains. Meta-domain analysis of γ-H2A.v enrichment in PR-Set7 depleted cells relative to control cells (blue) and replication timing values (orange) over late replicating domains. Each late replicating domain was divided into 50 bins and the average γ-H2A.v log2 difference between PR-Set7 depleted and control cells as well as replication timing values were plotted.

## DISCUSSION

Depletion of PR-Set7 in *Drosophila* Kc167 cells results in a decrease in H4K20 monomethylation, activation of an ATR-dependent checkpoint, and a cell cycle arrest in late S phase. The activation of the ATR-dependent checkpoint was accompanied by the accumulation of DNA damage. We found that PR-Set7 and H4K20 monomethylation of nascent histones were not required to complete the first S phase, but rather cells entering their second cell cycle failed to complete S phase likely due to the accumulation of unmodified histone H4K20 during the previous cell cycle. Unlike in mammalian cells, the accumulation of unmodified histone H4K20 did not impact origin activation. The failure to complete S phase is associated with the enrichment of γ-H2A.v in late replicating domains, suggesting that H4K20 methylation is critical in maintaining the integrity of late replicating sequences.

H4K20 is the most common histone target for post-translational modification. The vast majority (>80%) of H4K20 exists in the dimethyl state(7); therefore, every cell cycle the bulk of nascent histone H4 deposited during DNA replication needs to undergo monomethylation by PR-Set7 and subsequent di- or tri-methylation by Suv4-20. The conserved PIP-box mediating the PCNA and CRL4^Cdt2^ dependent degradation of PR-Set7 tightly couples PR-Set7 activity to the cell cycle. PR-Set7 globally monomethylates the nascent pool of H4K20 on lysine 20 in late S phase. Depletion of PR-Set7 in G1 and loss of late S phase specific H4K20 monomethylation had no impact on the progression through S phase in the first cell cycle, indicating that there is not an immediate “just in time” requirement for H4K20me1 to complete S phase or enter into mitosis. DNA damage and cell cycle arrest occur in the next cell cycle with the accumulation of nascent nucleosomes containing unmodified H4K20 throughout the genome (Fig 7).

**Figure 7.**
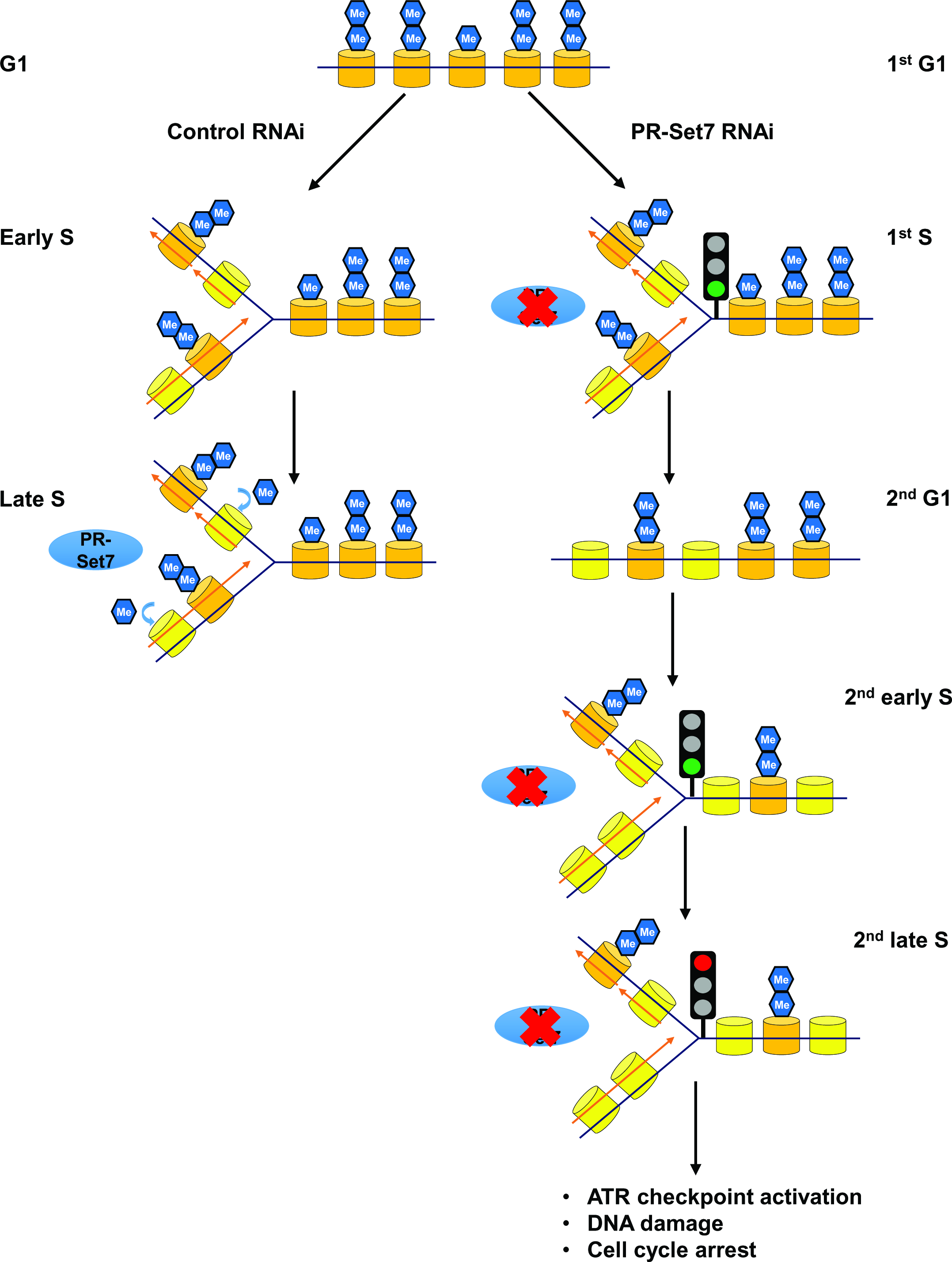
Model for the maintenance of *Drosophila* genome integrity by PR-Set7 in late replicating domains. The majority of parental histone H4 (orange) exists in the dimethylated state in G1. *Left branch (wild type):* PR-Set7 is absent in early S phase and nucleosomes containing nascent histone H4 unmethylated at H4K20 accumulate on newly synthesized DNA. PR-Set7 levels and activity are restored in late S phase and there is a global conversion of nascent nucleosomes from H4K20me0 to H4K20me1. Following S phase the monomethylated H4K20 is converted to H4K20me2. *Right branch (absence of PR-Set7):* In the absence of PR-Set7, the first S phase is completed and unmethylated nascent histone H4 (H4K20me0) is accumulated throughout the genome. In the subsequent S phase, early replicating domains are able to be copied; however, during late S phase there is an accumulation of DNA damage, activation of an ATR-dependent intra-S phase dependent checkpoint and cell cycle arrest. Presumably the presence of H4K20me0 in late replicating but not early replicating domains leads to replication fork stalling and collapse suggesting a critical role for PR-Set7 in maintaining the integrity of late replicating domains.

DNA damage resulting from loss of PR-Set7 and H4K20 methylation was restricted to late replicating regions of the genome, suggesting a defect in the DNA replication program at the end of S phase. A paucity of activated origins has been shown to lead to DNA damage, fragile sites, and increased mutations in human cells due to the increased distances replication forks must traverse(44-46). However, unlike in mammalian cells, deregulation of H4K20 methylation did not impair origin activation in *Drosophila.* In mammalian cells, tethering of PR-Set7 to the DNA promotes the locus-specific recruitment of replication factors and pre-RC assembly (presumably via increased levels of H4K20me2 and ORC stabilization)(11, 19, 20). However, given that almost all nucleosomes contain at least one histone H4 methylated at lysine 20, it is difficult to envision how tethering could lead to a greater local concentration of H4K20me2. It remains to be seen if the locus-specific effects observed at select origins in mammalian cells will represent a genome-wide paradigm(11).

Loss of PR-Set7 leads to a cell cycle arrest with nearly 4N DNA content. Depletion of the putative DNA damage checkpoint ATM did not relieve the cell cycle arrest. This may be due to the dispensable role of *Drosophila* ATM in maintaining the premitotic-checkpoint in response to DNA damage(47, 48). Instead, depletion of ATR activity relieved the checkpoint and cell cycle arrest suggesting that loss of PR-Set7 and H4K20 methylation activate the ATR-dependent intra-S phase checkpoint. The checkpoint is likely activated due to the accumulation of ssDNA and/or DSBs resulting from replication fork stalling and collapse (Reviewed in(49)). In the absence of the ATR-dependent checkpoint, the PR-Set7 depleted cells continued to cycle and we observed an attenuation of DNA damage signal (Fig S8). Presumably, the aneuploid tissue culture cells can accumulate modest DNA damage without immediate and catastrophic effects as previously reported(18). We also found that the pattern of γ-H2A.v marked DNA damage was broadly distributed across late replicating domains suggesting a model whereby the accumulation of unmodified H4K20 may render the chromatin recalcitrant to replication fork progression resulting in stochastic replication fork collapse.

Interestingly, the DNA damage associated with late replicating regions in the absence of PR-Set7 was restricted to late replicating euchromatic regions of the genome and not the pericentric heterochromatin (Fig S7). Why is *Drosophila* PR-Set7 and H4K20 methylation critical for the integrity of late euchromatic replicating domains? In contrast to early replicating domains which are enriched for activating chromatin PTMs and heterochromatic regions which are marked by methylated H3K9 or H3K27, late replicating regions of the euchromatic genome are generally gene-poor and depleted of most chromatin PTMs(41, 50). Thus, H4K20me1 may be a critical epigenetic mark to maintain proper chromatin architecture of euchromatic late replication domains (classified as "BLACK" chromatin by Filion et al.(50)). Consistent with this hypothesis, H4K20me1 is necessary to ensure proper chromatin compaction(18, 51). Loss of PR-Set7 leads to altered nuclear morphology; specifically, an enlarged nucleus due to abnormal chromatin decompaction(18). Together, these results suggest that normal chromosome compaction is a determinant of replisome stability and fork progression.

Alternatively, H4K20 methylation may be required for repairing and restarting spontaneous stalled replication forks which are more likely to be enriched in late replication regions (Reviewed in(52)). H4K20me2 has been implicated in interacting with 53BP1 in mammalian cells, which promotes the non-homologous end-joining (NHEJ) pathway to repair DNA DSBs and antagonizes the homologous recombination (HR) pathway factor BRCA1(53, 54). It is proposed that PR-Set7 deposits *de novo* monomethylation marks on H4K20 flanking DSBs, which provide substrates for sequential conversion to H4K20me2 by Suv4-20h1(55, 56). In mammalian U2OS cells, depletion of either an HR factor, Rad51, or an essential replication factor, Cdc45, eliminated the accumulation of γ-H2AX induced by PR-Set7 siRNA(14). These results suggest that PR-Set7 activity may promote NHEJ over HR during S phase. However, co-depletion of PR-Set7 and the *Drosophila* Rad51 or BRCA2 homolog had no impact on reducing the intensity of γ-H2A.v by immunofluorescence in our system (Fig S9). Also, loss of *Drosophila* Suv4-20 had little if any impact on the cell cycle progression of Kc167 cells (Fig 1B) or animal survival(57). This may reflect the variability of the replicative stress response pathways across species, or likely a less essential role of H4K20me2 in *Drosophila*, despite it being the most abundant form of H4K20(7).

PR-Set7 is essential for development. Loss of PR-Set7 in mice leads to embryonic lethality prior to the eight-cell stage(51); similarly, flies homozygous null for PR-Set7 die as third instar larvae (40). While it has been assumed that the essential function of the PR-Set7 methyltransferase is to monomethylate H4K20, it remains possible that it could target other critical proteins for methylation. Indeed, nonhistone targets of PR-Set7's catalytic activity have been identified in mammalian cells, including p53 (38) and PCNA(39), both of which are critical cell cycle regulators.

Recently, McKay and colleagues developed a novel histone replacement system to directly assess the function of specific histone lysine residues in *Drosophila* (25). Strikingly, more than half of the flies harboring a mutation of histone H4 lysine 20 to alanine (H4K20A), although sick, were able to survive into adulthood. H4K20A mutant flies also exhibited robust EdU incorporation in germline nurse and somatic follicle cells, which is consistent with our findings that origin activation is not impacted by loss of PR-Set7 activity. Despite the surprising viability of the mutant H4K20A flies, they did exhibit elevated DNA damage in proliferating larval wing discs (Fig 4) consistent with our hypothesis that methylation of H4K20 is a target of PR-Set7 in regulating genome integrity.

Despite the tight cell cycle-coupled regulation of PR-Set7, we found that depletion of PR-Set7 and loss of H4K20 monomethylation did not impair the first S phase but rather impacted the subsequent cell cycles. While we cannot completely rule out the possibility that it may take more than 24 hours for a PR-Set7 methylated non-histone protein to turn over, we suggest that cell cycle arrest in subsequent cell cycles is an epigenetic effect coupled to the accumulation of unmodified H4K20 on the chromatin. What is unclear, is precisely how much H4K20 methylation throughout the genome is required to maintain genomic stability? In the case of the H4K20A mutant flies, it remains possible that the incorporation of minimal amounts of the replication independent histone H4, His4r, (which was not mutated) may be sufficient to prevent the accumulation of DNA damage during late S phase. Finally, the majority of tissues derived from differentiated cells in the fly are polyploid resulting from developmentally programmed endocycles(58). These cells have under-replicated regions which frequently correspond with late replicating domains (59) and are de-sensitized to the DNA damage response pathway (60). Thus, H4K20 methylation may have a minimal impact on cells that are predisposed to fail to replicate and/or activate the DNA damage response in late replicating regions of the genome.

## ACKNOWLEDGEMENT

We thank members of the MacAlpine and Fox laboratories for suggestions and critical reading of the manuscript. We are grateful for bioinformatic advice and suggestions from Jeff Du and Jason A. Belsky.

### FUNDING

This work was supported by the American Cancer Society [RSG-11-048-01-DMC to D.M.M.] and the National Institutes of Health [T32-GM007092 to R.L.A., R01-DA036897 to R.J.D., and GM104097 to D.M.M.]. Funding for open access charge: National Institutes of Health.

---

**Figure S1.**
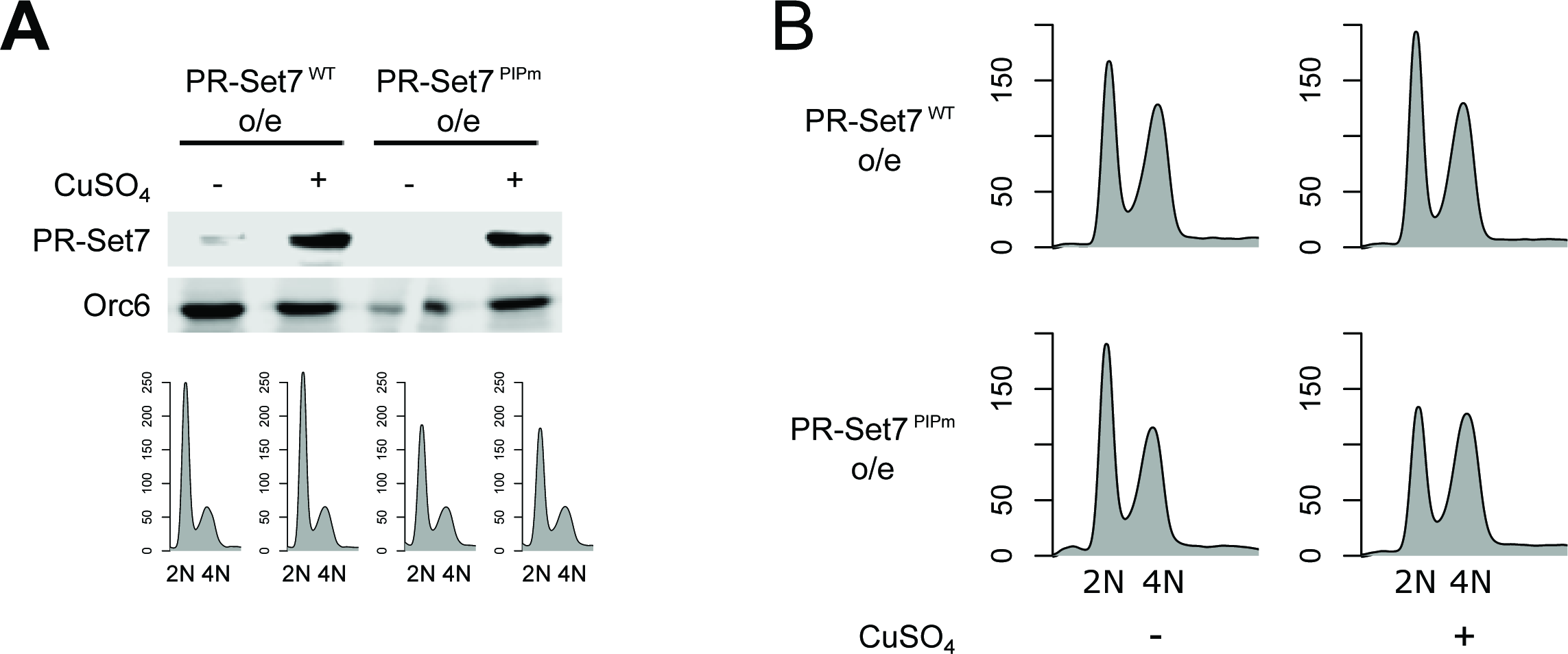
Overexpression of WT and PIP-mutant PR-Set7. (A) PR-Set7^WT^ or PR-Set7^PIPm^ cells were treated with HU and copper sulfate for 24 hours and harvested for the analysis of PR-Set7 levels by immunoblotting (Orc6 serves as a loading control). (B) Cell cycle profiles by flow cytometry after overexpression of asynchronous PR-Set7^WT^ or PR-Set7^PIPm^ for 72 hours.

**Figure S2.**
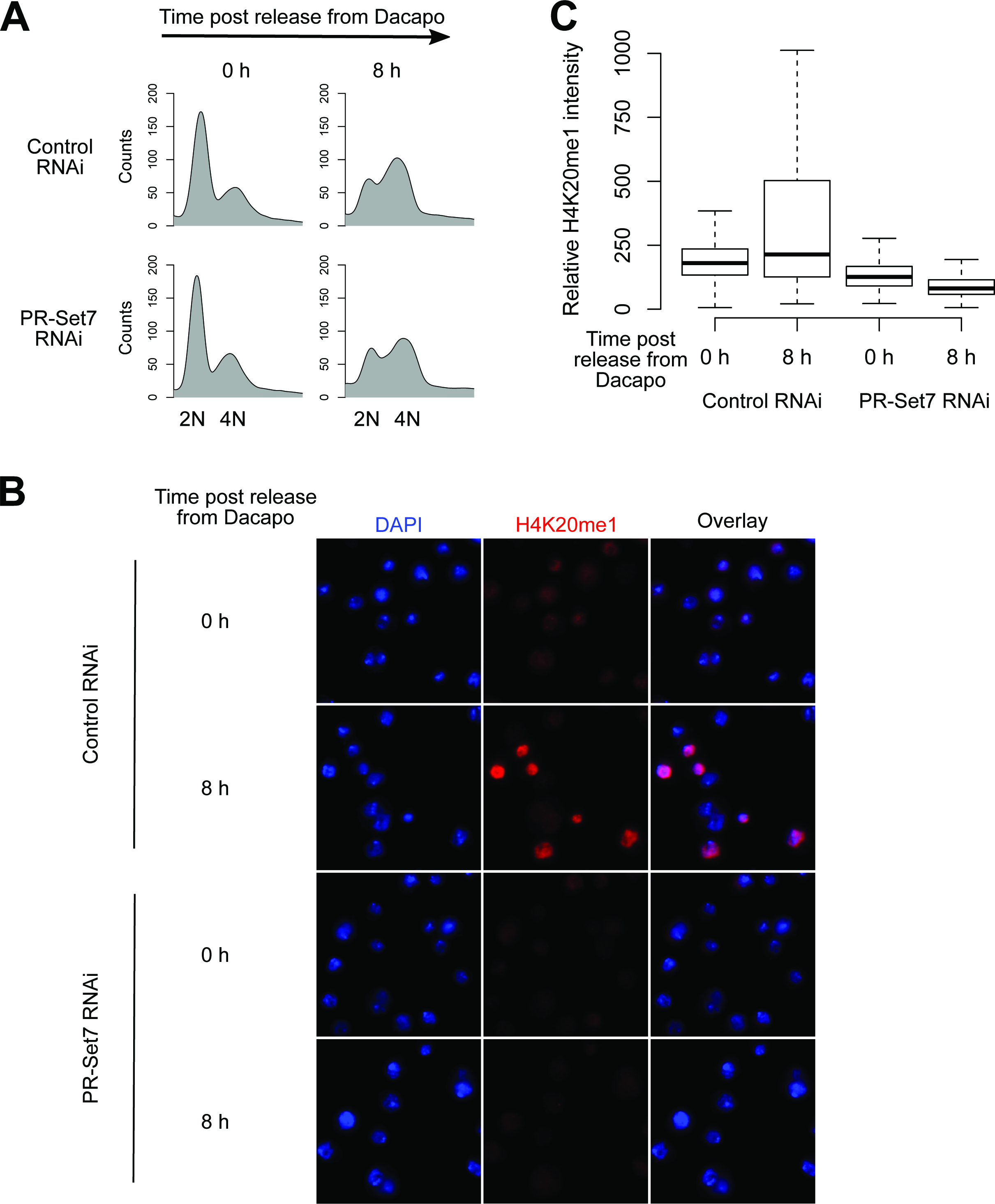
PR-Set7 depletion is sufficient to prevent the accumulation of H4K20me1 upon release from a Dacapo-induced G1 cell cycle arrest. (A) Control and PR-Set7 depleted cells enter late S/G2 within 8 hours following release from a Dacapo-induced G1 arrest. (B) RNAi-mediated depletion of PR-Set7 is sufficient to block the monomethylation of H4K20 in late S/G2 following release from the Dacapo induced cell cycle arrest. Immunofluorescence of H4K20me1 (red) in control and PR-Set7 depleted cells 8 hours following release into S phase. DNA is stained with DAPI (blue). (C) Boxplots resulting from the quantification of H4K20me1 intensity for each condition (n>180).

**Figure S3.**
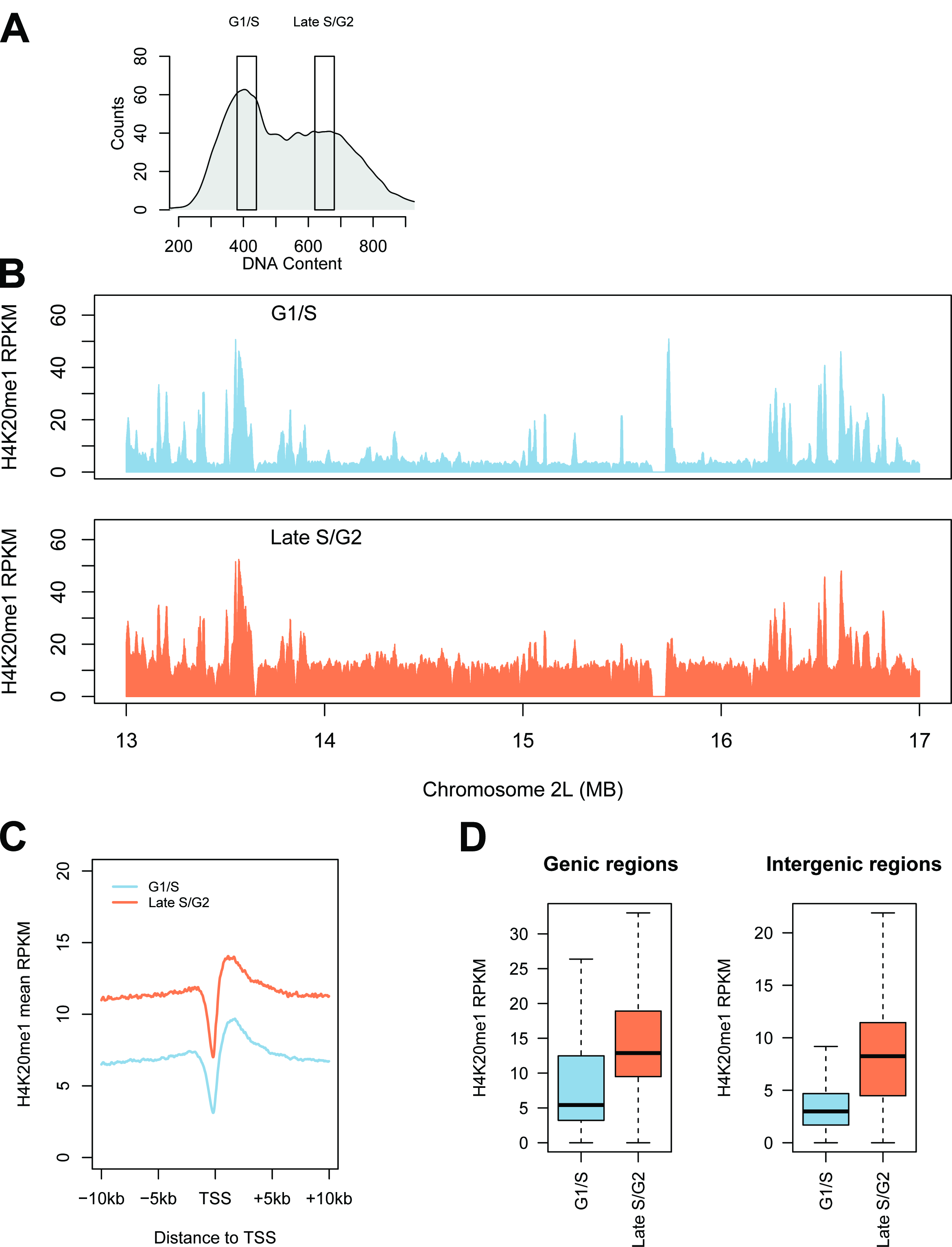
H4K20me1 levels are globally increased throughout the genome in late S phase. (A) Cell sorting. Hoechst 33342 stained cells were sorted into G1/S and late S/G2 populations based on DNA content. (B) Enrichment of H4K20me1 (RPKM) over a representative 4 MB region of chromosome 2L for cells enriched in G1/early S (blue) or late S/G2 (orange) of the cell cycle by cell sorting. (C) H4K20me1 is enriched at transcription start sites and gene bodies. Aggregate plots of H4K20me1 enrichment in G1/early S (blue) and late S/G2 (orange) cells relative to transcription start sites. (D) H4K20me1 levels increase in both genic and intergenic regions during late S phase. Boxplots representing the distribution of H4K20me1 RPKM within 13428 genic and 11430 intergenic regions (p<2.2×10^−16^).

**Figure S4.**
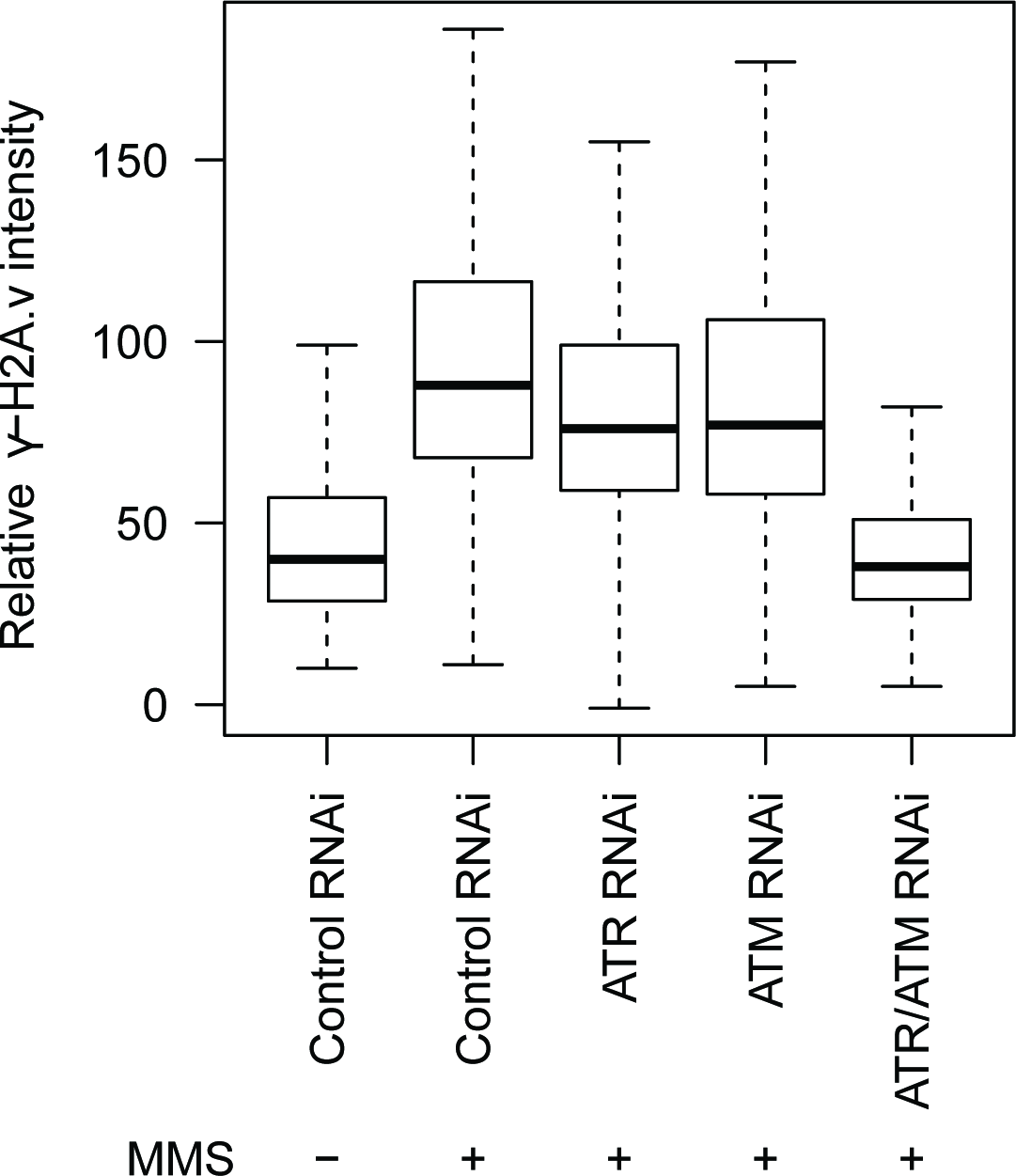
Verification of ATR and ATM RNAi efficacy. The cells were incubated with control, ATR, ATM, or both ATR and ATM dsRNA for 48 hours before 0.05% MMS was added and incubated for 2 hours (labeled as “MMS +”). The intensity of γ-H2A.v phosphorylation (γ-H2A.v) in response to MMS treatment was measured by immunofluorescence microscopy and depicted with boxplots (n > 250).

**Figure S5.**
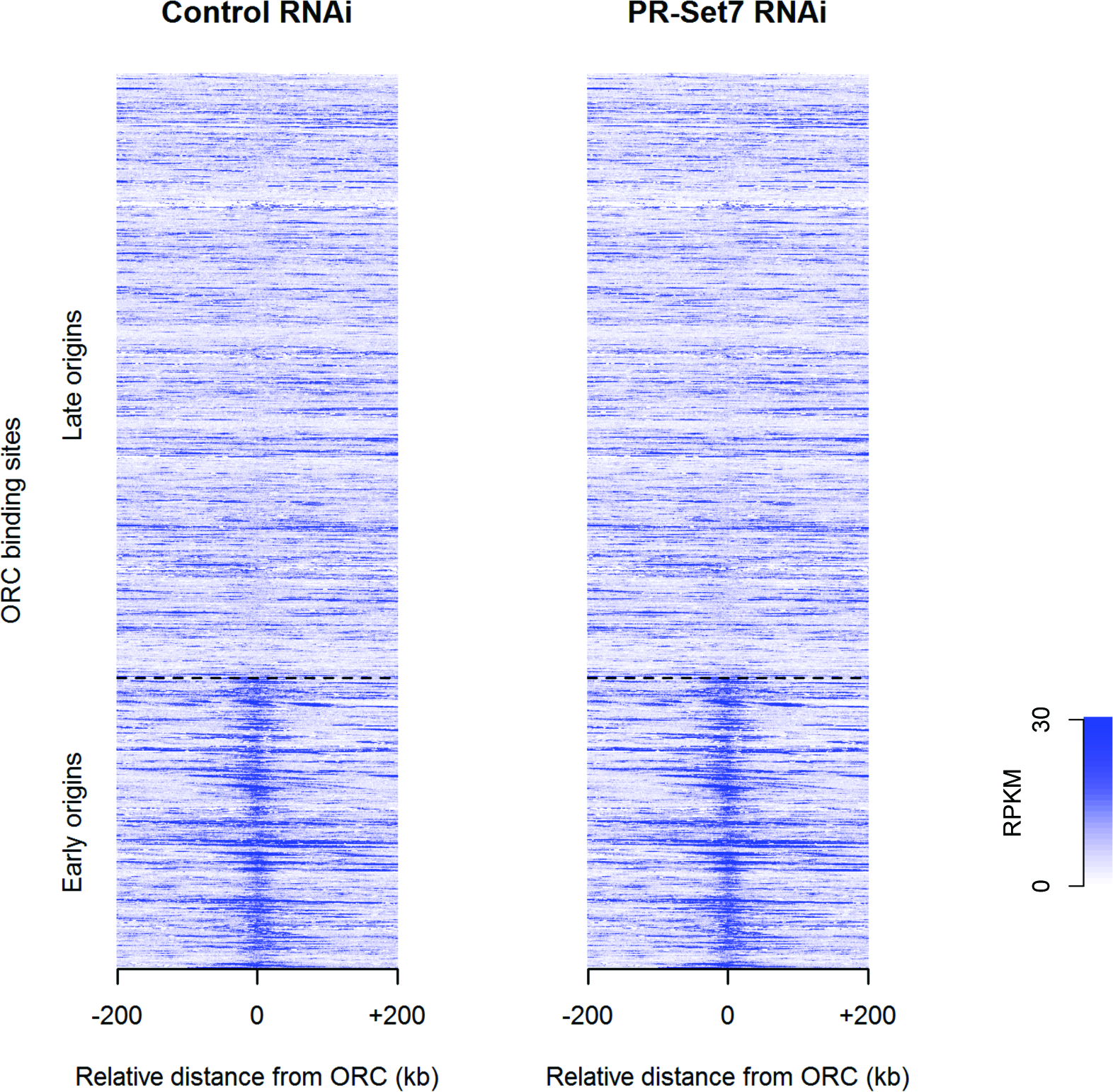
BrdU incorporation in HU arrested cells is enriched at ORC binding sites. Heatmap of BrdU incorporation (measured as RPKM) in relation to 5159 ORC binding sites (41). Below the dashed line are the 1677 ORC sites at early origins and above the dashed line are ORC sites at late origins. The center of ORC peak is labeled as position 0.

**Figure S6.**
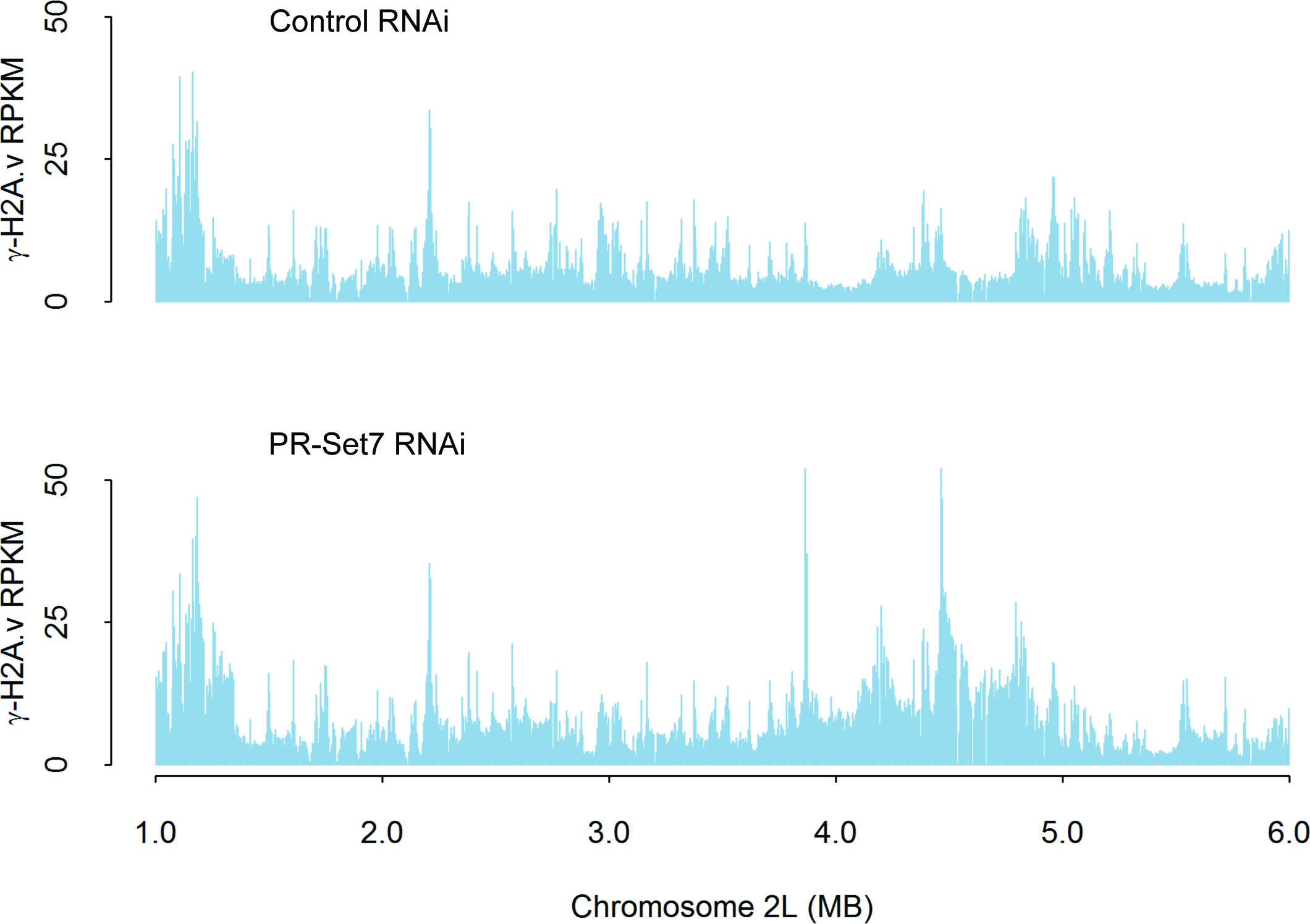
Genome-wide distribution of γ-H2A.v for control and PR-Set7 depleted cells. ChIP-seq analysis of the genome-wide distribution of γ-H2A.v for control and PR-Set7 depleted cells (RPKM).

**Figure S7.**
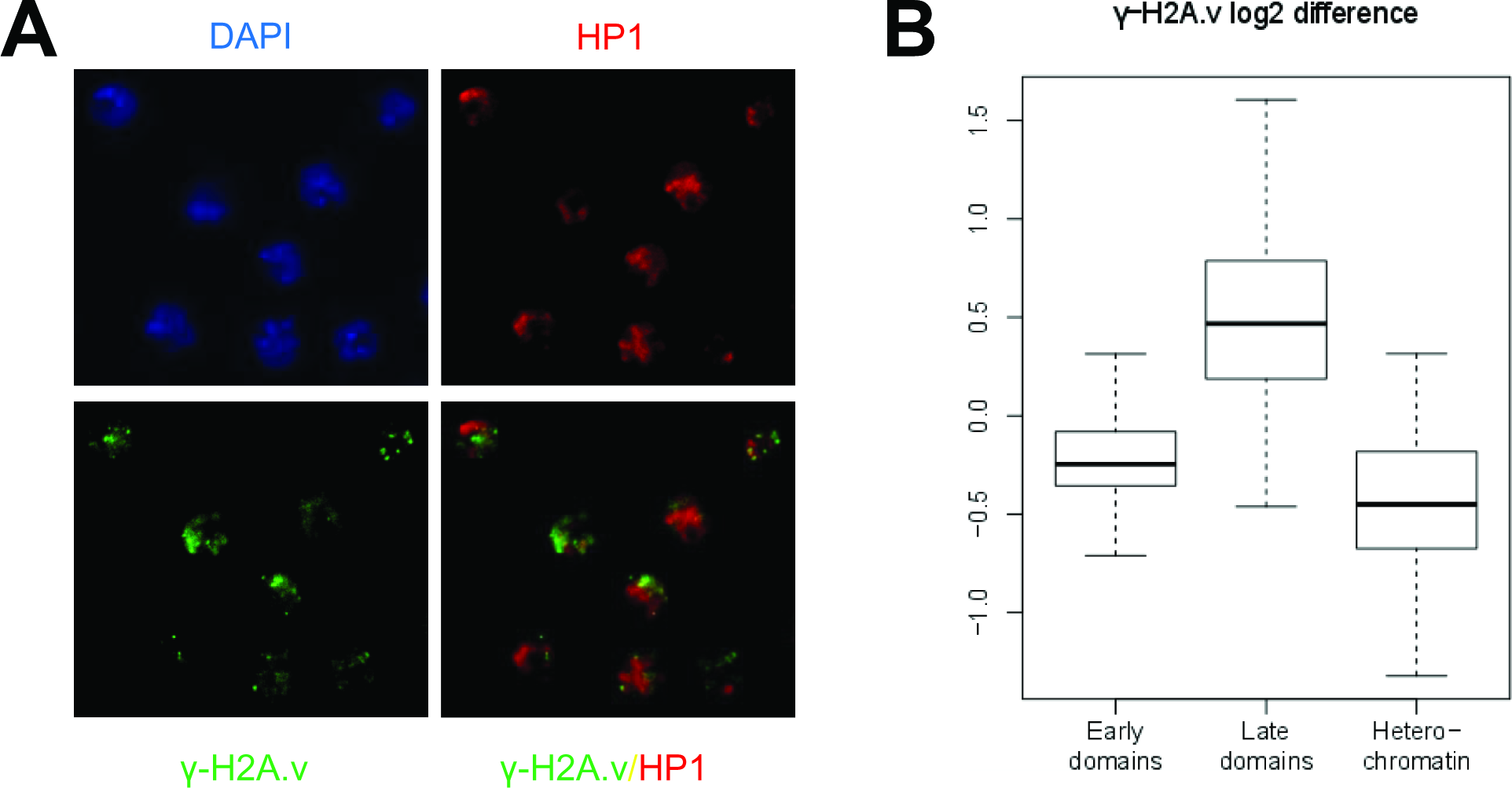
Loss of PR-Set7 induced DNA damage does not specifically localize to the heterochromatin. (A) Asynchronous cells treated with PR-Set7 RNAi for 72 hours before immunofluorescence staining of HP1 (red; Abcam ab24726, 1:250) and γ-H2A.v (green). DNA is counterstained with DAPI (blue). (B) γ-H2A.v ChIP-seq datasets analyzed in Fig 6 were re-aligned to include repetitive pericentric heterochromatic regions (Bowtie parameter: -M 1). The γ-H2A.v log2 difference was calculated as in Fig 6. Boxplots of the log2 difference in γ-H2A.v signal between control and PR-Set7 RNAi depleted cells in early euchromatic replicating domains, late euchromatic replicating domains, and heterochromatic regions are shown.

**Figure S8.**
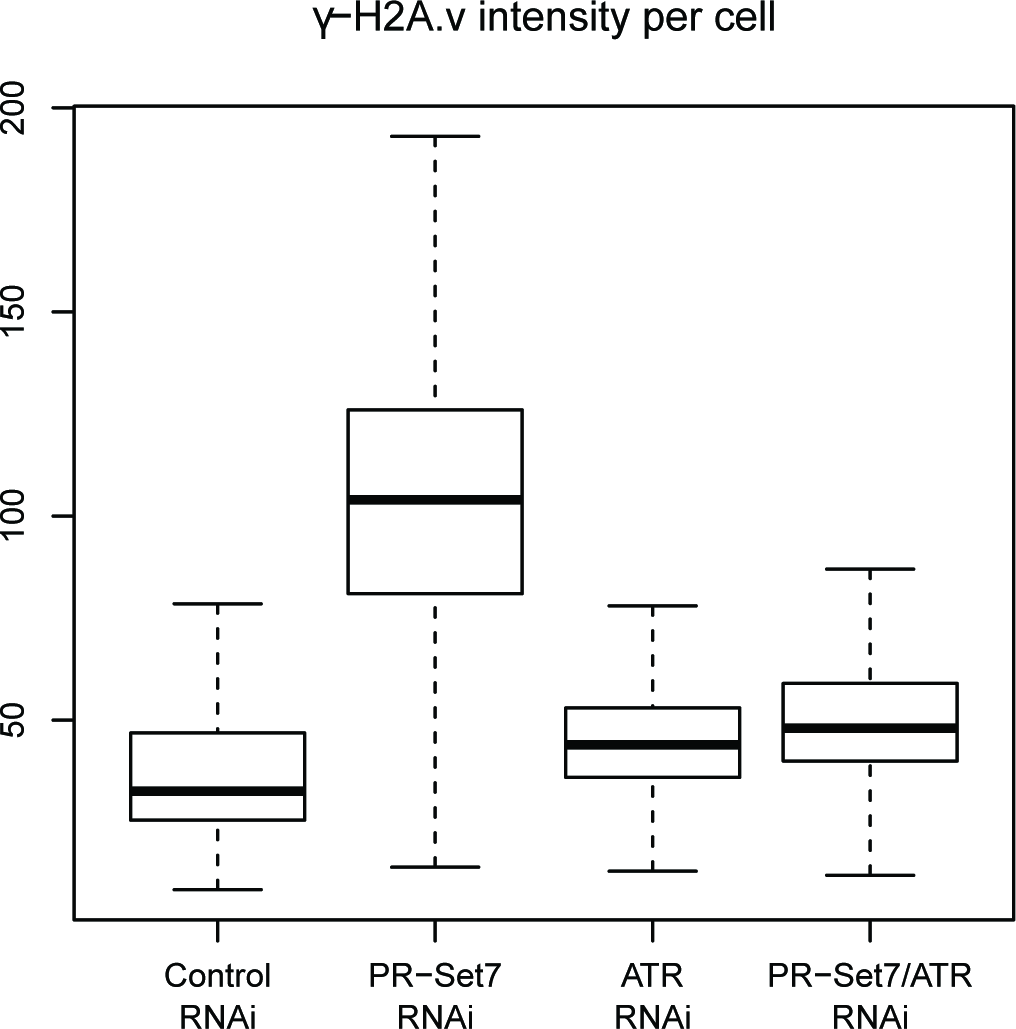
ATR co-depletion reduces γ-H2A.v caused by loss of PR-Set7. Asynchronous cells were treated with control, PR-Set7, ATR, and PR-Set7/ATR RNAi for 72 hours and harvested for immunofluorescence staining with antibodies against γ-H2A.v. The γ-H2A.v intensity was determined and depicted in boxplots for each condition (n > 1100).

**Figure S9.**
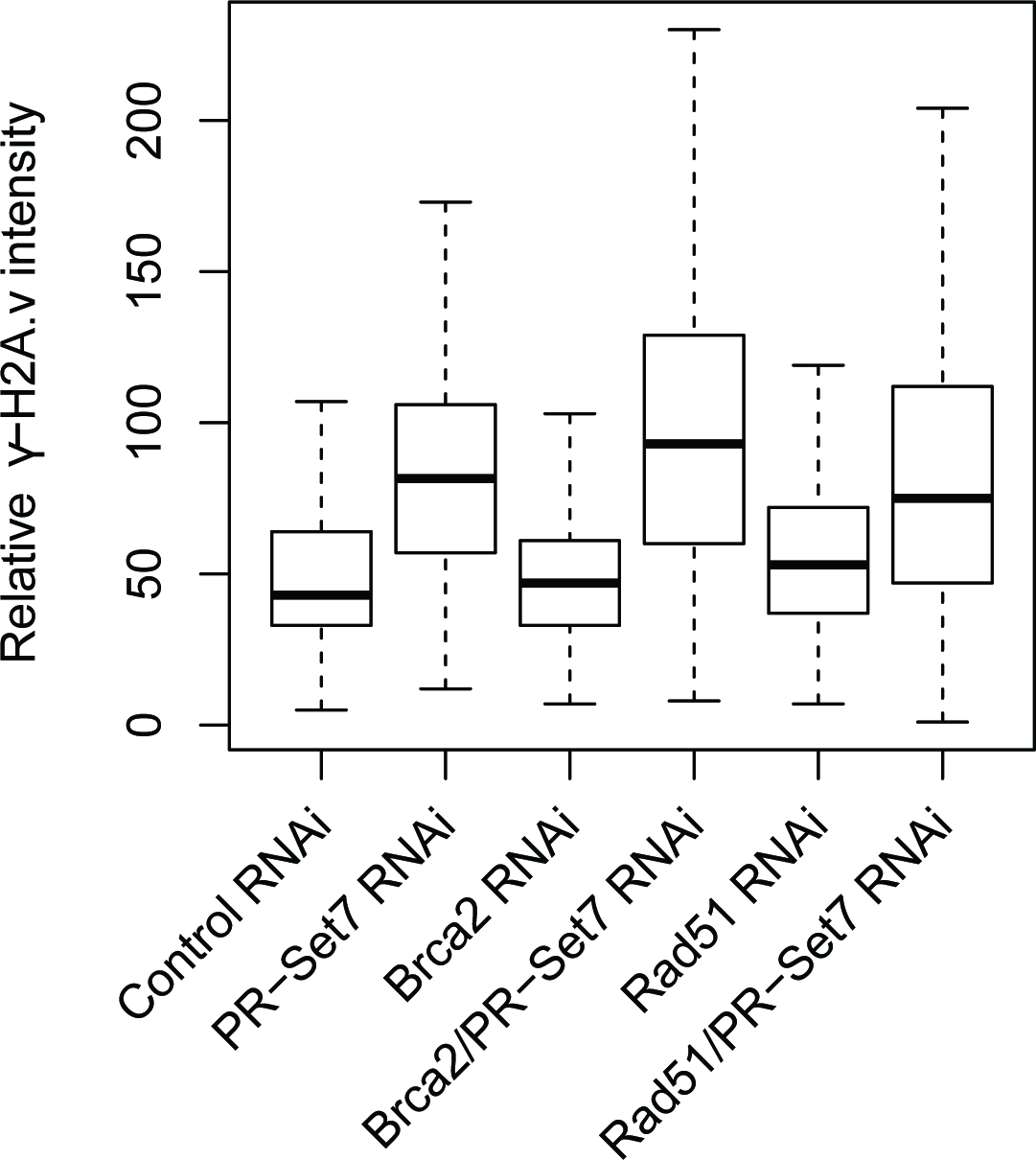
Disruption of the homologous recombination (HR) pathway does not reduce DNA damage resulting from loss of PR-Set7. Cells were treated with control or PR-Set7 RNAi for 72 hours before harvest and immunofluorescence staining of γ-H2A.v. The immunofluorescence intensity of γ-H2A.v under each condition was depicted with boxplots (n > 250).

**Table S1.** Chromosomal positions of late replicating domains and γ-H2A.v domains identified by HMM.

| | Late replicating domains | | | γ-H2A.v domains | | |
| --- | --- | --- | --- | --- | --- | --- |
|  | Chr | Start | End | Chr | Start | End |
| 1 | 2L | 627002 | 813000 | 2L | 580000 | 810000 |
| 2 | 2L | 1005002 | 1048000 | 2L | 900000 | 1080000 |
| 3 | 2L | 1258002 | 1350000 | 2L | 1170000 | 1590000 |
| 4 | 2L | 2406002 | 2648000 | 2L | 1730000 | 1910000 |
| 5 | 2L | 3839002 | 4200000 | 2L | 2220000 | 2730000 |
| 6 | 2L | 4268002 | 4326000 | 2L | 3620000 | 4880000 |
| 7 | 2L | 4457002 | 4793000 | 2L | 5520000 | 5850000 |
| 8 | 2L | 6148002 | 6294000 | 2L | 6100000 | 6340000 |
| 9 | 2L | 8806002 | 8864000 | 2L | 7500000 | 7800000 |
| 10 | 2L | 10069002 | 10138000 | 2L | 10510000 | 10950000 |
| 11 | 2L | 10601002 | 10842000 | 2L | 11240000 | 12000000 |
| 12 | 2L | 11318002 | 11787000 | 2L | 12100000 | 12440000 |
| 13 | 2L | 11831002 | 12002000 | 2L | 13900000 | 16270000 |
| 14 | 2L | 12135002 | 12362000 | 2L | 16850000 | 18540000 |
| 15 | 2L | 14087002 | 14178000 | 2L | 19190000 | 19360000 |
| 16 | 2L | 14419002 | 14486000 | 2L | 19570000 | 19750000 |
| 17 | 2L | 14540002 | 15011000 | 2L | 19820000 | 20070000 |
| 18 | 2L | 15090002 | 15239000 | 2L | 20440000 | 20650000 |
| 19 | 2L | 15281002 | 15654000 | 2L | 21750000 | 22070000 |
| 20 | 2L | 15779002 | 16221000 | 2R | 2940000 | 3130000 |
| 21 | 2L | 16962002 | 17339000 | 2R | 4610000 | 4780000 |
| 22 | 2L | 17517002 | 18105000 | 2R | 9450000 | 9710000 |
| 23 | 2L | 18181002 | 18409000 | 2R | 11270000 | 11410000 |
| 24 | 2L | 19835002 | 20110000 | 2R | 12230000 | 12430000 |
| 25 | 2L | 20223002 | 20372000 | 2R | 13030000 | 13280000 |
| 26 | 2L | 20488002 | 20573000 | 2R | 14730000 | 15040000 |
| 27 | 2L | 21795002 | 22112000 | 2R | 15360000 | 16150000 |
| 28 | 2R | 1683002 | 1791000 | 2R | 16210000 | 16790000 |
| 29 | 2R | 2396002 | 2454000 | 2R | 17630000 | 17970000 |
| 30 | 2R | 2957002 | 3086000 | 2R | 18570000 | 18700000 |
| 31 | 2R | 4185002 | 4322000 | 2R | 18820000 | 19260000 |
| 32 | 2R | 4622002 | 4799000 | 3L | 1870000 | 2390000 |
| 33 | 2R | 4837002 | 4972000 | 3L | 2840000 | 3050000 |
| 34 | 2R | 5533002 | 5724000 | 3L | 3600000 | 3820000 |
| 35 | 2R | 6187002 | 6296000 | 3L | 4410000 | 5750000 |
| 36 | 2R | 6844002 | 6899000 | 3L | 5880000 | 6120000 |
| 37 | 2R | 7357002 | 7423000 | 3L | 6210000 | 6920000 |
| 38 | 2R | 7605002 | 7723000 | 3L | 7010000 | 7130000 |
| 39 | 2R | 7953002 | 8008000 | 3L | 7390000 | 7730000 |
| 40 | 2R | 9524002 | 9679000 | 3L | 9130000 | 9340000 |
| 41 | 2R | 10534002 | 10641000 | 3L | 9850000 | 10640000 |
| 42 | 2R | 10910002 | 10965000 | 3L | 11790000 | 12040000 |
| 43 | 2R | 11280002 | 11369000 | 3L | 13030000 | 13260000 |
| 44 | 2R | 11457002 | 11670000 | 3L | 13360000 | 13920000 |
| 45 | 2R | 12245002 | 12425000 | 3L | 14180000 | 14480000 |
| 46 | 2R | 13828002 | 13889000 | 3L | 14780000 | 14980000 |
| 47 | 2R | 14057002 | 14157000 | 3L | 15300000 | 15490000 |
| 48 | 2R | 14750002 | 15022000 | 3L | 15690000 | 15950000 |
| 49 | 2R | 15414002 | 15580000 | 3L | 16230000 | 16350000 |
| 50 | 2R | 15667002 | 16094000 | 3L | 17020000 | 17430000 |
| 51 | 2R | 16244002 | 16491000 | 3L | 18100000 | 18670000 |
| 52 | 2R | 16552002 | 16869000 | 3L | 19080000 | 19390000 |
| 53 | 2R | 17696002 | 17930000 | 3L | 20530000 | 20710000 |
| 54 | 2R | 18120002 | 18181000 | 3L | 21670000 | 21820000 |
| 55 | 2R | 18710002 | 18761000 | 3L | 21940000 | 22260000 |
| 56 | 2R | 18851002 | 18929000 | 3L | 22400000 | 22720000 |
| 57 | 2R | 18972002 | 19237000 | 3R | 1610000 | 2200000 |
| 58 | 2R | 19322002 | 19435000 | 3R | 2270000 | 2580000 |
| 59 | 2R | 20278002 | 20386000 | 3R | 3030000 | 3740000 |
| 60 | 3L | 1981002 | 2147000 | 3R | 4170000 | 4360000 |
| 61 | 3L | 2203002 | 2350000 | 3R | 6700000 | 7030000 |
| 62 | 3L | 2686002 | 2731000 | 3R | 7830000 | 8070000 |
| 63 | 3L | 2862002 | 3064000 | 3R | 8540000 | 8820000 |
| 64 | 3L | 3640002 | 3760000 | 3R | 8860000 | 9150000 |
| 65 | 3L | 3955002 | 4008000 | 3R | 9210000 | 9400000 |
| 66 | 3L | 4720002 | 5094000 | 3R | 10710000 | 10920000 |
| 67 | 3L | 5381002 | 5569000 | 3R | 12940000 | 13440000 |
| 68 | 3L | 5838002 | 5887000 | 3R | 13550000 | 13880000 |
| 69 | 3L | 5947002 | 6068000 | 3R | 14500000 | 15590000 |
| 70 | 3L | 6293002 | 6878000 | 3R | 15700000 | 16110000 |
| 71 | 3L | 7017002 | 7113000 | 3R | 16140000 | 16370000 |
| 72 | 3L | 7156002 | 7234000 | 3R | 17890000 | 18200000 |
| 73 | 3L | 7446002 | 7716000 | 3R | 18610000 | 18870000 |
| 74 | 3L | 9204002 | 9293000 | 3R | 19170000 | 19550000 |
| 75 | 3L | 9921002 | 10203000 | 3R | 21120000 | 21340000 |
| 76 | 3L | 10508002 | 10582000 | 3R | 21970000 | 22630000 |
| 77 | 3L | 11835002 | 12045000 | 3R | 23100000 | 24630000 |
| 78 | 3L | 12586002 | 12663000 | 3R | 24750000 | 24860000 |
| 79 | 3L | 13060002 | 13187000 | 3R | 25100000 | 25480000 |
| 80 | 3L | 13573002 | 13799000 | 3R | 26390000 | 26570000 |
| 81 | 3L | 14310002 | 14370000 | 3R | 26850000 | 27030000 |
| 82 | 3L | 14846002 | 14962000 | 3R | 27270000 | 27410000 |
| 83 | 3L | 15358002 | 15418000 | 3R | 27630000 | 27770000 |
| 84 | 3L | 15740002 | 15786000 | X | 2660000 | 2910000 |
| 85 | 3L | 15851002 | 15908000 | X | 3070000 | 3180000 |
| 86 | 3L | 16247002 | 16330000 | X | 4590000 | 5310000 |
| 87 | 3L | 17087002 | 17437000 | X | 6250000 | 6430000 |
| 88 | 3L | 18201002 | 18584000 | X | 7210000 | 7410000 |
| 89 | 3L | 19166002 | 19226000 | X | 9240000 | 9390000 |
| 90 | 3L | 19294002 | 19580000 | X | 9580000 | 10130000 |
| 91 | 3L | 20584002 | 20758000 | X | 10770000 | 11130000 |
| 92 | 3L | 20862002 | 20970000 | X | 11910000 | 12400000 |
| 93 | 3L | 21056002 | 21125000 | X | 12670000 | 13020000 |
| 94 | 3L | 21644002 | 21805000 | X | 13310000 | 13470000 |
| 95 | 3L | 21977002 | 22246000 | X | 13880000 | 14920000 |
| 96 | 3L | 22453002 | 22702000 | X | 15020000 | 15190000 |
| 97 | 3L | 24280002 | 24398000 | X | 17860000 | 18140000 |
| 98 | 3R | 1667002 | 2164000 | X | 19800000 | 20280000 |
| 99 | 3R | 2332002 | 2586000 | X | 20370000 | 20910000 |
| 100 | 3R | 2650002 | 2839000 | X | 21320000 | 21430000 |
| 101 | 3R | 3059002 | 3300000 | X | 21620000 | 21700000 |
| 102 | 3R | 3400002 | 3594000 | X | 21910000 | 22440000 |
| 103 | 3R | 3635002 | 3710000 |  |  |  |
| 104 | 3R | 6260002 | 6576000 |  |  |  |
| 105 | 3R | 6732002 | 6982000 |  |  |  |
| 106 | 3R | 7107002 | 7346000 |  |  |  |
| 107 | 3R | 7923002 | 8153000 |  |  |  |
| 108 | 3R | 8361002 | 8424000 |  |  |  |
| 109 | 3R | 8600002 | 8800000 |  |  |  |
| 110 | 3R | 8918002 | 9068000 |  |  |  |
| 111 | 3R | 9207002 | 9408000 |  |  |  |
| 112 | 3R | 9667002 | 9767000 |  |  |  |
| 113 | 3R | 10755002 | 10937000 |  |  |  |
| 114 | 3R | 11566002 | 11609000 |  |  |  |
| 115 | 3R | 12317002 | 12449000 |  |  |  |
| 116 | 3R | 12491002 | 12785000 |  |  |  |
| 117 | 3R | 12983002 | 13191000 |  |  |  |
| 118 | 3R | 13271002 | 13337000 |  |  |  |
| 119 | 3R | 13648002 | 13966000 |  |  |  |
| 120 | 3R | 15055002 | 15413000 |  |  |  |
| 121 | 3R | 15867002 | 16066000 |  |  |  |
| 122 | 3R | 16176002 | 16367000 |  |  |  |
| 123 | 3R | 17910002 | 18226000 |  |  |  |
| 124 | 3R | 18634002 | 18819000 |  |  |  |
| 125 | 3R | 19224002 | 19471000 |  |  |  |
| 126 | 3R | 20204002 | 20304000 |  |  |  |
| 127 | 3R | 20776002 | 20826000 |  |  |  |
| 128 | 3R | 20971002 | 21039000 |  |  |  |
| 129 | 3R | 21187002 | 21263000 |  |  |  |
| 130 | 3R | 22132002 | 22244000 |  |  |  |
| 131 | 3R | 22390002 | 22522000 |  |  |  |
| 132 | 3R | 23177002 | 23375000 |  |  |  |
| 133 | 3R | 23428002 | 23735000 |  |  |  |
| 134 | 3R | 23900002 | 24048000 |  |  |  |
| 135 | 3R | 24195002 | 24338000 |  |  |  |
| 136 | 3R | 24481002 | 24610000 |  |  |  |
| 137 | 3R | 24720002 | 24828000 |  |  |  |
| 138 | 3R | 25210002 | 25529000 |  |  |  |
| 139 | 3R | 26316002 | 26505000 |  |  |  |
| 140 | 3R | 26910002 | 27013000 |  |  |  |
| 141 | X | 2747002 | 2791000 |  |  |  |
| 142 | X | 3910002 | 3981000 |  |  |  |
| 143 | X | 4301002 | 4409000 |  |  |  |
| 144 | X | 4652002 | 4817000 |  |  |  |
| 145 | X | 4892002 | 5170000 |  |  |  |
| 146 | X | 5900002 | 6067000 |  |  |  |
| 147 | X | 7284002 | 7375000 |  |  |  |
| 148 | X | 8190002 | 8255000 |  |  |  |
| 149 | X | 9273002 | 9387000 |  |  |  |
| 150 | X | 9630002 | 10122000 |  |  |  |
| 151 | X | 10768002 | 10955000 |  |  |  |
| 152 | X | 11994002 | 12289000 |  |  |  |
| 153 | X | 13375002 | 13448000 |  |  |  |
| 154 | X | 13954002 | 14078000 |  |  |  |
| 155 | X | 14204002 | 14532000 |  |  |  |
| 156 | X | 14588002 | 14654000 |  |  |  |
| 157 | X | 15058002 | 15157000 |  |  |  |
| 158 | X | 17172002 | 17370000 |  |  |  |
| 159 | X | 17811002 | 17949000 |  |  |  |
| 160 | X | 18028002 | 18144000 |  |  |  |
| 161 | X | 18859002 | 18915000 |  |  |  |
| 162 | X | 19852002 | 20037000 |  |  |  |
| 163 | X | 20101002 | 20249000 |  |  |  |
| 164 | X | 20423002 | 20877000 |  |  |  |
| 165 | X | 21335002 | 21456000 |  |  |  |
| 166 | X | 22028002 | 22211000 |  |  |  |

**Table S2.**
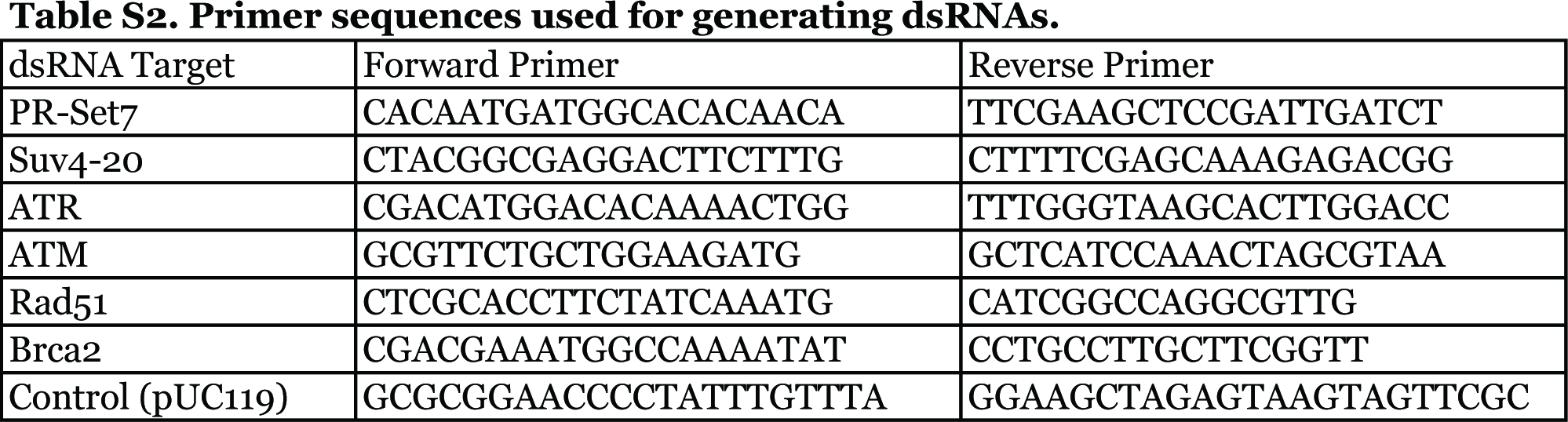
Primer sequences used for generating dsRNAs.

**Table S3.** List of biological replicates for H4K20me1 ChIP-seq, origin activity, and γ-H2A.v ChIP-seq genomic assays and the pearson correlations between individual replicate datasets.

| Experiment | Sample | Replicate # | Pearson correlation |
| --- | --- | --- | --- |
| H4K20me1 ChIP-seq | HU arrest | 1 | 0.949 |
|  | sorted G1/S | 1 |  |
|  | HU release | 1 | 0.901 |
|  | sorted late S/G2 | 1 |  |
| Early origin | control RNAi | 1 | #1, #2: 0.898 |
|  |  | 2 | #2, #3: 0.921 |
|  |  | 3 | #1, #3: 0.958 |
|  | PR-Set7 RNAi | 1 | 0.847 |
|  |  | 2 |  |
|  | ATR RNAi | 1 | 0.726 |
|  |  | 2 |  |
|  | ATR/PR-Set7 RNAi | 1 | 0.876 |
|  |  | 2 |  |
|  | mock treatment | 1 |  |
|  | Suv4-20 RNAi | 1 | 0.966* |
|  | PR-Set7/Suv4-20 RNAi | 1 |  |
|  | PR-Set7 o/e | 1 | 0.953* |
|  | PR-Set7 o/e + Suv4-20 RNAi | 1 |  |
| γ-H2A.v ChIP-seq | control RNAi | 1 | 0.974 |
|  |  | 2 |  |
|  | PR-Set7 RNAi | 1 | 0.975 |
|  |  | 2 |  |

\* Samples are genetically redundant, so they could serve as biological replicates.

